# Enforced specificity of an animal symbiosis

**DOI:** 10.1101/2024.08.04.606548

**Authors:** Julian M. Wagner, Jason H. Wong, Jocelyn G. Millar, Enes Haxhimali, Adrian Brückner, Thomas H. Naragon, James Q. Boedicker, Joseph Parker

**Affiliations:** Division of Biology and Biological Engineering, California Institute of Technology, Pasadena, CA, USA; Department of Physics and Astronomy, University of Southern California, Los Angeles, CA, USA; Department of Entomology, University of California, Riverside, CA, USA; Division of Chemistry and Chemical Engineering, California Institute of Technology, Pasadena, CA, USA

## Abstract

Insect diversification has been catalyzed by widespread specialization on novel hosts—a process underlying exceptional radiations of phytophagous beetles, lepidopterans, parasitoid wasps, and inordinate lineages of symbionts, predators and other trophic specialists. The strict fidelity of many such interspecies associations is posited to hinge on sensory tuning to host-derived cues, a model supported by studies of neural function in host-specific model species. Here, we investigated the sensory basis of symbiotic interactions between a myrmecophile rove beetle and its single, natural host ant species. We show that host cues trigger analogous behaviors in both ant and symbiont. Cuticular hydrocarbons—the ant’s nestmate recognition pheromones—elicit partner recognition by the beetle and execution of ant grooming behavior, integrating the beetle into the colony via chemical mimicry. The beetle also follows host trail pheromones, permitting inter-colony dispersal. Remarkably, the rove beetle also performs its symbiotic behaviors with non-host ants separated by ∼95 million years, and shows minimal preference for its natural host over non-host ants. Experimentally validated agent-based modeling supports a scenario in which specificity is enforced by physiological constraints on symbiont dispersal, and negative fitness interactions with alternative hosts, rather than via sensory tuning. Enforced specificity may be a pervasive mechanism of host range restriction of specialists embedded within host niches. Chance realization of latent compatibilities with alternative hosts may facilitate host switching, enabling deep-time persistence of obligately symbiotic lineages.

---

Ecological specialization permeates all domains of life^1,2^—the outcome of an evolutionary narrowing of niche breadth stemming from a diversity of context- and taxon-specific drivers^1,3,4^. Despite the pervasiveness of specialization, the biological mechanisms that functionally constrain organisms to a single or limited spectrum of hosts remain incompletely known for most specialist lineages^5,6^. Among insects, specialization is rife, manifesting in clades of trophic^7–10^, mutualistic^11,12^ or parasitic^13–18^ specialists that target highly restricted host ranges, often obligately so. Attraction to host-derived sensory cues is typical of insect specialists (e.g. ^19–26^), and is often posited to underlie the tight fidelity of these associations^7,21,27–33^. Efficient sensory processing of host cues has been shown to confer adaptive value, enabling specialists to more effectively recognize and exploit partner species, whilst simultaneously constraining the specialist’s interaction space^27,34–36^. Neural-based models of host specificity are supported by studies of the insect nervous system, where specialists display enhanced sensitivity to host cues at the sensory periphery (e.g.^37–40^) and in central brain circuits^41–43^, as well as greater anatomical investment in brain regions associated with host cue transduction^38,44–46^. Yet, explicit tests of whether sensory tuning suffices to explain patterns of host specificity in nature are scarce; in addition to sensory information, ecological forces including pressure from natural enemies^47,48^, limitations in dispersal capacity or host encounter probability^49–52^, and the potential for locating conspecifics^53,54^ have been proposed to shape the realized host range. A major conundrum is the widespread ability of specialists to switch to phylogenetically divergent hosts over evolutionary time—a paradox given the obligate lifestyles and extreme selectivity of many specialists^6,55–57^. Consequently, the mechanisms, both neural and ecological, that govern host associations over ecological and evolutionary timescales remain unresolved.

Myrmecophiles are symbiotic organisms adapted for life inside ant societies, and represent an archetype of extreme ecological specialization^58^. Approximately 100,000 such species are estimated to exist^59^, including many phenotypically elaborate taxa that have evolved sophisticated behavioral and chemical strategies to infiltrate and parasitically exploit host colonies^58,60–63^. Myrmecophilous associations are often socially complex, obligate, and highly host specific to single ant species^60,64–72^. Rapid mortality of some myrmecophiles is observed on removal from the host colony environment^64^. Myrmecophiles therefore provide a paradigm for unravelling the mechanisms underlying obligate, symbiotic dependencies on specific host organisms. Evidence indicates that myrmecophiles can be strongly attracted to host ants, performing behaviors such as ant grooming^60,68,73–79^, phoretic attachment to ant bodies^73,80–83^, mouth-to-mouth feeding (trophallaxis)^68,84–89^ and navigating ant foraging trails^90–93^. To date, however, knowledge of the sensory cues that myrmecophiles use to find, recognize and interact with ants is scarce. The basis for the extraordinary fidelity of myrmecophile-ant relationships remains unknown, but must be reconciled with the counterintuitive observation of host promiscuity of many myrmecophile taxa over evolutionary time. Host switching is prominent across ancient, speciose clades of obligate myrmecophiles, and has likely been central to the persistence and diversification of these organisms^87,94–98^.

Here, we exploit the biology of myrmecophiles to test the forces shaping the interaction space of extreme ecological specialists. By harnessing a naturally occurring ant-myrmecophile relationship, we have been able to reconstitute a behaviorally complex animal symbiosis in the laboratory, study it using automated, quantitative methods, and experimentally deconstruct it to expose the symbiont’s mechanisms of host finding, host recognition, and host specificity. The product of this work is a theoretically and empirically supported model in which a symbiont’s natural host range is an emergent property of the agency of both symbiont and host organisms. By yielding a solution to the host switching paradox, our findings move towards a unified understanding of how the same forces govern patterns of symbiotic association over ecological and evolutionary timescales.

## The myrmecophile *Sceptobius*

Three ant species of the genus *Liometopum* (Formicidae: Dolichoderinae) are widespread in southwestern North America, forming vast colonies of ∼10^6^ workers that forage across hundreds of square meters of habitat^99^. Each *Liometopum* species plays host to one of three myrmecophile species of the rove beetle genus *Sceptobius* (Staphylinidae). The beetles are hypothesized to have co-speciated with *Liometopum*^100–102^ (**Fig. 1A**), each beetle being host-specific despite partially sympatric ranges of the three ant species (**Fig. 1B**). Of these host-symbiont pairings, *Sceptobius lativentris* and *Liometopum occidentale* are abundant across southern California, including field sites in the Angeles National Forest (CA: LA County). *S. lativentris* exhibits strict partner fidelity in nature, having been recorded exclusively from colonies of *L. occidentale*^101^. The beetles are tightly integrated into the host society and strongly attracted to ants. When placed with a worker ant, *Sceptobius* will climb onto its body, clasping the ant’s antenna in its mandibles (**Fig. 1C; Video S1a, b**). Secured to the ant in this way, the beetle proceeds to repeatedly “groom” the worker’s body surface with its tarsi, alternating with rubbing its tarsi over its own body. Grooming behavior functions to transfer cuticular hydrocarbons (CHCs)—the ant’s nestmate recognition pheromones, which form a waxy coating on workers’ bodies^103–105^—onto the beetle’s cuticle. Via this social interaction, the beetle achieves perfect chemical mimicry, its CHC profile identically matching that of the ant, enabling it to gain acceptance into its host colony (**Fig. 1D**).

**Figure 1:**
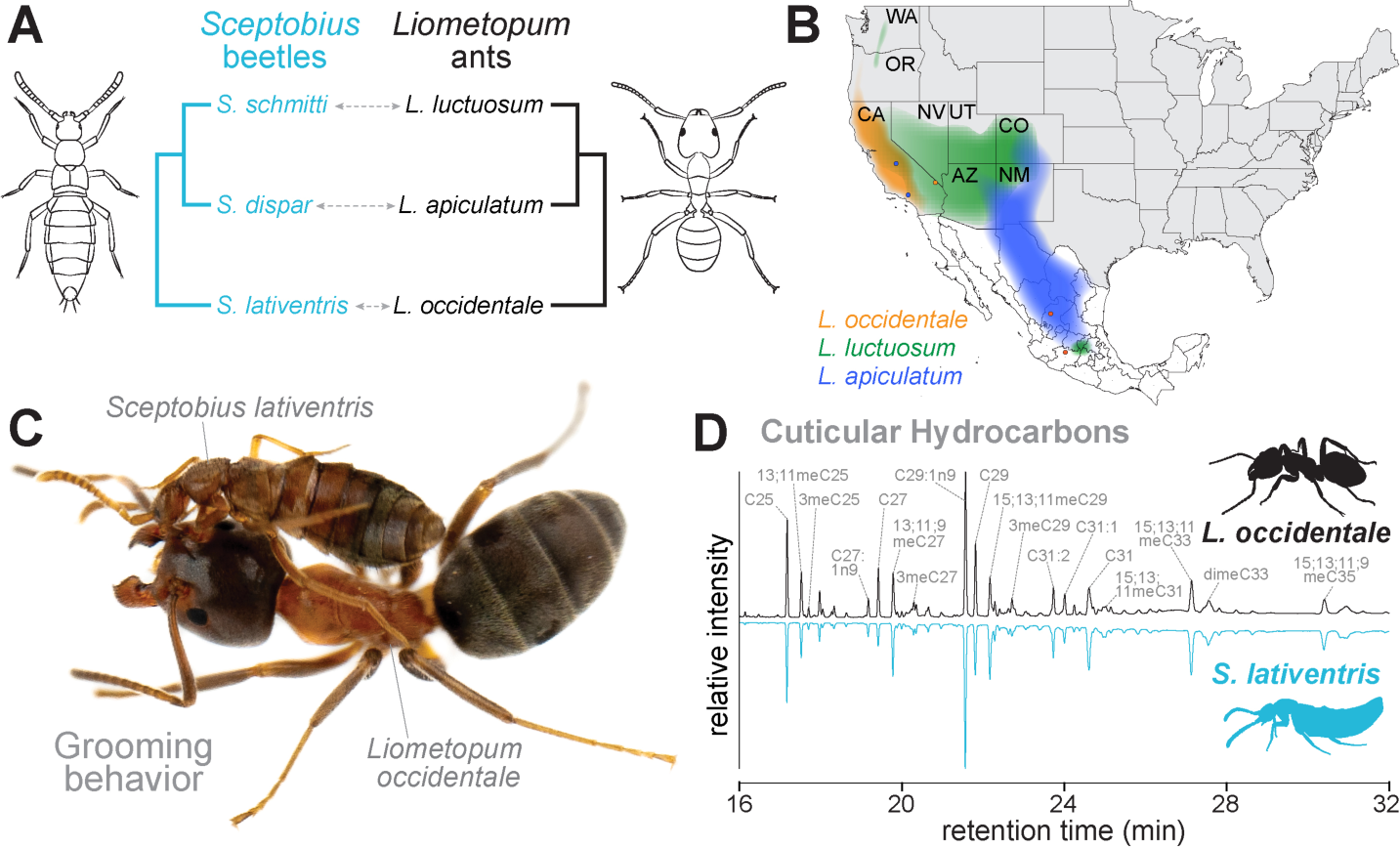
A model ant-myrmecophile system. (**A**) Three symbiotic *Sceptobius* species associate with corresponding *Liometopum* host species. (**B**) Ranges of *Liometopum* species in North America, showing partially sympatric ranges. (**C**) *S. lativentris* grooming a *L. occdentale* worker. Grooming functions to acquire the ant’s CHC profile, leading to perfect chemical mimicry by the symbiont. (**D**) Gas chromatograh traces of *L. occidentale* ant (black) and *S. lativentris* beetle (blue), with identities of CHC peaks indicated.

Importantly, the ant’s CHCs also provide a waxy barrier that safeguards against desiccation^106^. A chronic, physically close association with host ants is thus essential for the survival of the myrmecophile; *Sceptobius* may spend over half of its adult life grooming ants^101^. The beetles are unable to live away from their hosts, die rapidly when isolated from colonies, and have evolved to be flightless^100^ (see **File S1** for details of *Sceptobius* life history). Like other myrmecophiles, however, *Sceptobius* can disperse from nests by navigating along actively used *L. occidentale* foraging trails, which traverse the forest floor. Such deep assimilation into its host ant’s biology is presumably the outcome of long-term evolution within the *Liometopum* colony niche. We asked what mechanisms functionally tie this obligate symbiont to its single, specific partner species.

## Sensory control of myrmecophile host recognition

We have found that the symbiotic biology of *Sceptobius* can be reconstituted in the laboratory. The beetle readily performs highly stereotyped interactions with ants in experimental contexts, allowing us to examine their sensory control, and test the sufficiency of these behaviors in explaining the beetle’s natural host specificity. We constructed a behavioral platform to quantitatively study the beetle’s grooming behavior, comprising a multiplexed array of interaction arenas, illuminated with infrared light to eliminate visual stimuli (**Fig 2A; Fig. S1A, B**). Into these arenas we placed single pairs of beetles and host ants. By training a deep-learning neural network^107^ to follow keypoints on both insects, we tracked host and symbiont movement in arenas over periods of 2 hours (**Fig. 2A**, **Fig. S1C**). During these trials, *Sceptobius* climbed onto the ant and performed repeated grooming bouts, which could be easily classified by clustering of beetle and ant keypoints within 3 mm for at least 30 seconds, (**Fig. 2B; Video S2**). Individual grooming bouts varied in duration, but often lasted many minutes, and sometimes for over 1 hr (**Fig. 2G**).

**Figure 2:**
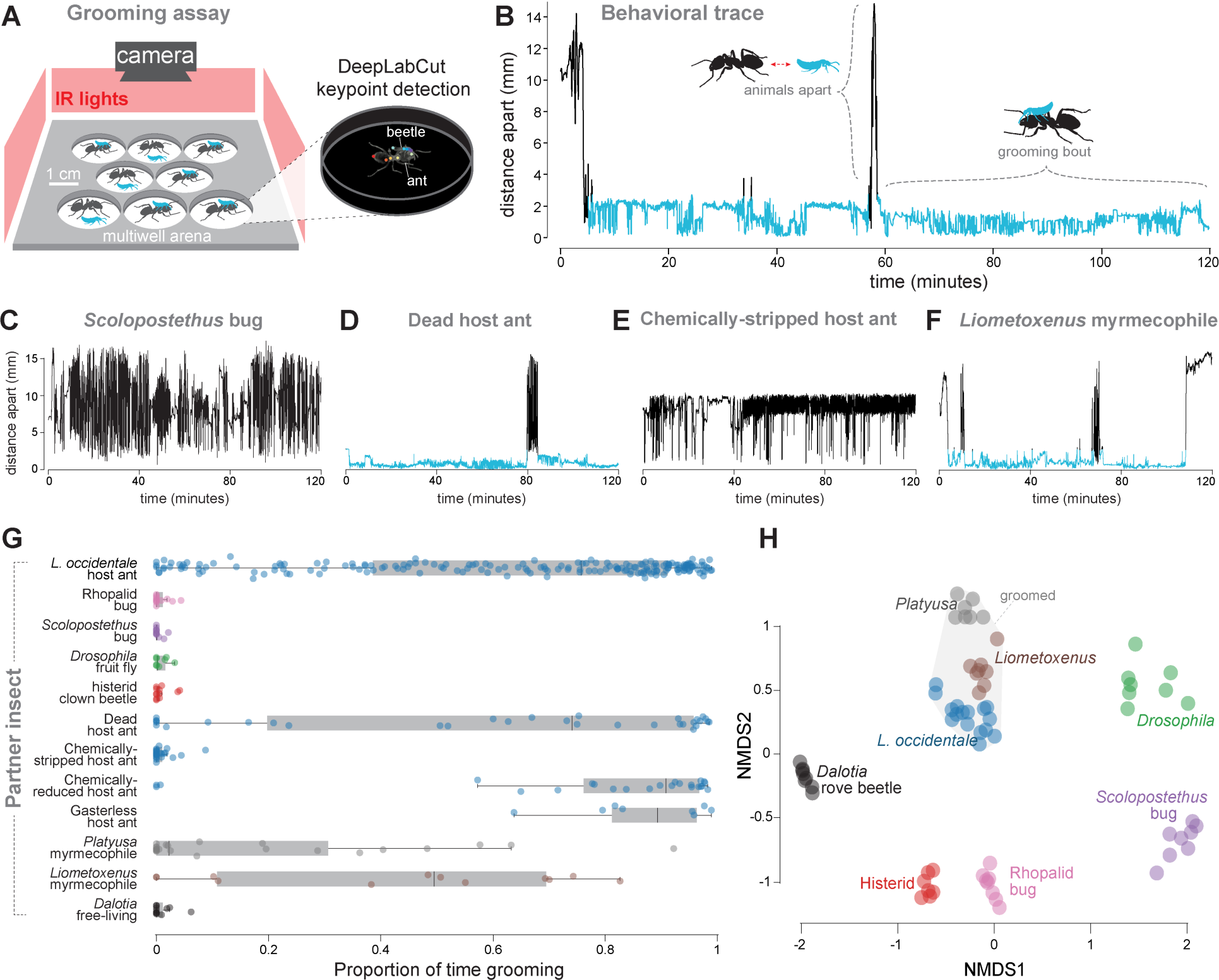
CHCs are host recognition cues. (**A**) Multi-arena behavioral platform with multiple animal position with DeepLabCut to quantify grooming behavior. (**B**) Representative 2-hour behavioral trace. Blue stretches indicate grooming bouts where beetles and ants converge for ≥30 seconds; black indicates periods of non-grooming. (**C**) *Sceptobius* does not groom a hemipteran bug. (**D**) *Sceptobius* grooms dead *L. occidentale* worker ants. (**E**) *Sceptobius* does not groom dead, chemically-stripped *L. occidentale* workers. (**F**) *Sceptobius* grooms other myrmecophiles that mimic *L. occidentale* CHCs. (**G**) Summary of grooming times during two-hour experiments. (**H**) NMDS plot of CHC profiles from insect species assayed for grooming. Insects that *Sceptobius* grooms cluster closely in CHC chemical space relative to non-groomed insects.

We explored the basis for the strong physical attraction of *Sceptobius* to its *L. occidentale* host ant. First, we substituted non-ant insects of the approximate same size and shape as *L. occidentale* into arenas with *Sceptobius*. On introduction of a hemipteran bug (*Scolopostethus* sp.) that is commonly found in leaf litter surrounding *L. occidentale* colonies^108^, no behavioral attraction from *Sceptobius* was observed (**Fig. 2C, G; Video S3**), implying that the beetle can distinguish its natural host from a non-ant insect. Identical results were obtained when *Sceptobius* was permitted to interact with other non-ants, including another hemipteran species, as well as fruit flies and histerid beetles; all species were ignored by *Sceptobius* (**Fig. 2G**). We hypothesized that *Sceptobius* recognizes its host based on chemosensory information on the ant body. Indeed, *Sceptobius* is attracted to and will groom dead *L. occidentale* worker ants, demonstrating that the beetle does not recognize kinematic features of its host (**Fig. 2D, G; Video S4**). Conversely, when the dead host is washed repeatedly in hexane to strip it of external chemical secretions, the grooming interaction is abolished, indicating that other features of the ant body, such as cuticle microsculpture, do not release grooming (**Fig. 2E, G; Video S5**). If the ant is hexane-washed to the point of strongly decreasing, but not completely removing, external chemical secretions, however, ant attraction and grooming remain intact (**Fig. 2G**). We conclude that *Sceptobius* is highly sensitive to chemicals on the ant body surface.

We further defined which types of ant compound elicit grooming. In hexane extracts of crude body washes of *L. occidentale* workers, we find three major compound classes: CHCs (the ant’s nestmate recognition cues), 6-methyl-5-hepten-2-one (sulcatone; a volatile compound emitted as an alarm pheromone^109^), and iridoids—a class of compound shown to function as trail pheromones in related dolichoderine ant species^110,111^. The sulcatone and iridoids are excreted by an abdominal pygidial gland^112^. Accordingly, crude hexane extracts of *L. occidentale* with gasters removed possess only CHCs, without detectable sulcatone and iridoids (**Fig. S2A; Video S6**). Nevertheless, we observe that *Sceptobius* grooms gasterless *L. occidentale* equivalently to intact ants (**Fig. 2G**; **Fig. S2B**), implying that CHCs—and not iridoids or sulcatone—are the relevant cues that elicit ant grooming. To unequivocally confirm that CHCs are the relevant host recognition cues, we took an unusual approach. We have found that two, phylogenetically distantly related myrmecophile rove beetle genera, *Platyusa* (Aleocharinae: Lomechusini) and *Liometoxenus* (Aleocharinae Oxypodini) have, in addition to *Sceptobius*, convergently evolved to target colonies of *L. occidentale*. Both *Platyusa* and *Liometoxenus* have evolved to accurately chemically mimic the CHCs of *L. occidentale*, displaying the same set of hydrocarbon compounds at similar ratios on their bodies as their ant hosts. These myrmecophiles cluster closely with the ant in chemical space relative to all the other non-ant insects we tested (**Fig. 2H**). Remarkably, when placed into arenas with either of these myrmecophile rove beetles, *Sceptobius* mounted and groomed them, much like it behaves towards *L. occidentale* (**Fig. 2F**; **Fig. 2G; Video S7; Video S8**). Conversely, *Sceptobius* did not groom a free-living rove beetle, *Dalotia*^113^ (Aleocharinae: Athetini) (**Fig. 2G**), CHCs of which are dissimilar to those of *L. occidentale* (**Fig. 2H**)^114^. We conclude that *Sceptobius* recognizes its *L. occidentale* host based on its host’s CHC profile. Detection of host ant CHCs triggers a social behavioral grooming program by which *Sceptobius* achieves chemical mimicry, integrating the beetle into the colony.

## Iridoid trail pheromones mediate host finding and dispersal

In addition to CHC-based host recognition, *Sceptobius* employs a mode of long-range movement that further ties it to its host. We commonly observe beetles walking along the extensive networks of *L. occidentale* foraging trails in the Angeles National Forest. Limited aggression and significant trail connectivity exist between distant *L. occidentale* colonies^115^. Trail following may permit the flightless, desiccation-prone beetle to disperse between colonies while remaining physically close to worker ants. We investigated the sensory basis of trail following by permitting an *L. occidentale* colony to forage through a large arena, into which were placed irregularly shaped obstacles (**Fig. 3A, Fig. S3A**). We tracked cumulative ant density over 12 hours to map a trail formed through the obstacles (**Fig. 3B, Fig. S3B**). We then removed the ants and obstacles and introduced a single worker ant into the vacant area (**Fig. S3C**). The ant’s movement corresponded closely to the region of highest ant density, confirming that the foraging ant colony left behind a robust chemical trail (**Fig. 3B, E; Fig. S3D**). Strikingly, when a *Sceptobius* beetle was introduced into the vacant arena, its movement also matched the shape of the *L. occidentale* trail near-perfectly (**Fig. 3C, E; Fig. S3E**). By contrast, neither movement of a free-living *Dalotia* rove beetle (**Fig. S3F**), or the movement of *Sceptobius* in an empty trail-free arena (**Fig. 3D, E**), showed any correspondence with ant trail shape. We deduce that *Sceptobius* has an evolved ability to follow chemical trails laid by its host ant.

**Figure 3:**
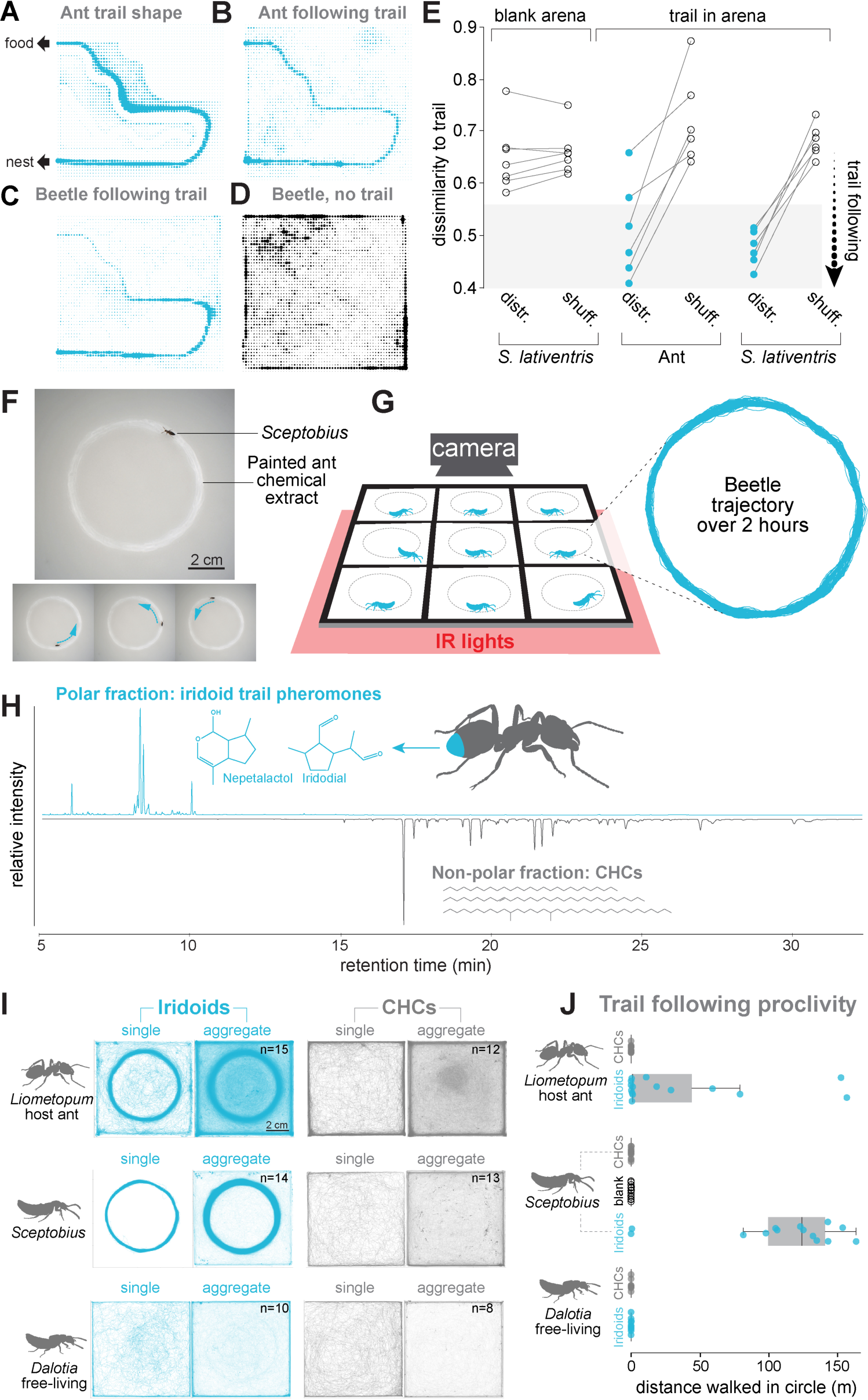
Evolution of iridoid trail following enables host finding. (**A-D**) Binned density plots of animal movement within a foraging arena. (**A**) Cumulative *L. occidentale* colony movement in arena shows trail formation around obstacles. Movement of single worker ant (**B**) or *Sceptobius* (**C**) in vacant arena following removal of colony and obstacles reveals high accuracy trail following behavior. (**D**) *Sceptobius* walks largely around arena perimeter in a host ant trail-free arena. (**E**) Trail following accuracy. Using a dissimilarity measure derived from the Bhattacharyya distance, movement of *S. lativentris* and the ant correspond closely to the trail distribution (‘distr.’); randomly shuffling beetle movement traces abolishes the close match to the trail distribution, indicating that random movement cannot account for the correspondence of beetle movement with ant trail (‘shuff.’), whereas beetle movement in an empty arena had an equally negligible fit to trail shape as a randomly shuffled movement trace. (**F**) *Sceptobius* follows ant pheromones painted onto glass. (**G**) Multiplexed behavioral arena to monitor trail following. (**H**) Fractionation of ant pheromones into polar and non-polar compounds yields iridoids (iridodial and nepetalactol) and CHCs, respectively. (**I**) Individual examples and aggregate trajectories of *L. occidentale* ants, *Sceptobius* beetles and free-living *Dalotia* beetles in arenas with painted iridoids or CHCs. Both worker ants and *Sceptobius* closely followed only the iridoid fraction, while *Dalotia* followed neither chemical fraction (**H**) Quantification of trail following distances during two-hour trials.

We found that when crude ant chemical extracts are painted in a ring-shape on a ground-glass arena floor, *Sceptobius* will follow the circular trail, often for many revolutions (tens to hundreds of meters) (**Fig. 3F, Video S9**). We exploited this assay to identify which *L. occidentale* compounds function as trail pheromones for both host ant and myrmecophile. By fractionating crude ant chemical extracts, we recovered a nonpolar portion containing the full complement of CHCs, and a polar fraction comprising a series of stereoisomers of two iridoids: iridodial and nepetalactol (**Fig. 3H**). Using a multiplexed, glass-floored arena, we quantified insect movement around circular trails of painted CHC- or iridoid-fraction over 2 hours (**Fig. 3G, Fig. S4A, B**). Both *L. occidentale* worker ants and the myrmecophile *Sceptobius* exclusively followed iridoid trails, confirming that these compounds are the trail pheromones (**Fig. 3I, J, Video S10-S13**). Again, the free-living beetle *Dalotia* failed to follow trails of either compound class (**Fig. 3I, J, Video S14**).

## Ant chemical cues do not mediate host specificity

Our findings demonstrate that *S. lativentris* has evolved to eavesdrop on two major components of ant communication and interprets them in a manner analogous to that of its host ant. The beetle uses the ant’s nestmate recognition cues—CHCs—as host recognition cues; additionally, the beetle follows the ant’s iridoid foraging trails for probable dispersal and host finding. Via these mechanisms, *Sceptobius* maintains a close association with its host. We hypothesized that these same chemical cues may mediate the natural specificity of *S. lativentris* to its single, *L. occidentale* host ant species. As demonstrated above, *Sceptobius* did not interact with insects lacking the requisite CHC profile, consistent with models in which sensitivity to host-derived cues underlies partner specificity of ecological specialists. We therefore employed our behavioral assays to assess whether *L. occidentale* chemical cues were the sole releasers of the beetle’s symbiotic behaviors. To our surprise, despite the absolute specificity of the *S. lativentris-L. occidentale* association in nature, *S. lativentris* is profoundly promiscuous in the laboratory. We observed that the beetle robustly performed grooming behavior with both *Liometopum* sister ant species, *L. luctuosum* and *L. apiculatum* (**Fig. 4A, B, Video S15**). Pushing the promiscuity further, we tested phylogenetically divergent ants, and found *Sceptobius* would groom ants from the subfamilies Myrmicinae (*Veromessor* and *Pogonomyrmex*) and Formicinae (*Formica*), which diverged from *Liometopum* approximately 95–100 million years ago^116^ (**Fig. 4A, C, Video S16**). Not only does the beetle recognize non-host ants and perform its symbiotic grooming behavior with them, but in each case, grooming shifts the beetle’s CHC profile to almost perfectly match that of the non-host ant (**Fig. 4D**). Using *L. luctuosum* as an experimental non-host ant, we quantified how grooming causes the beetles to acquire a new host ant species’ identity. We found a time-dependent shift in CHC profile, with complete chemical integration with non-hosts after only a day (**Fig. 4E**). Once integrated, we saw long-term survival of *S. lativentris* in experimental colonies of non-host ants, which can rescue the normal, rapid mortality of beetles when removed from colonies of their natural *L. occidentale* host (**Fig. 4F**).

**Figure 4:**
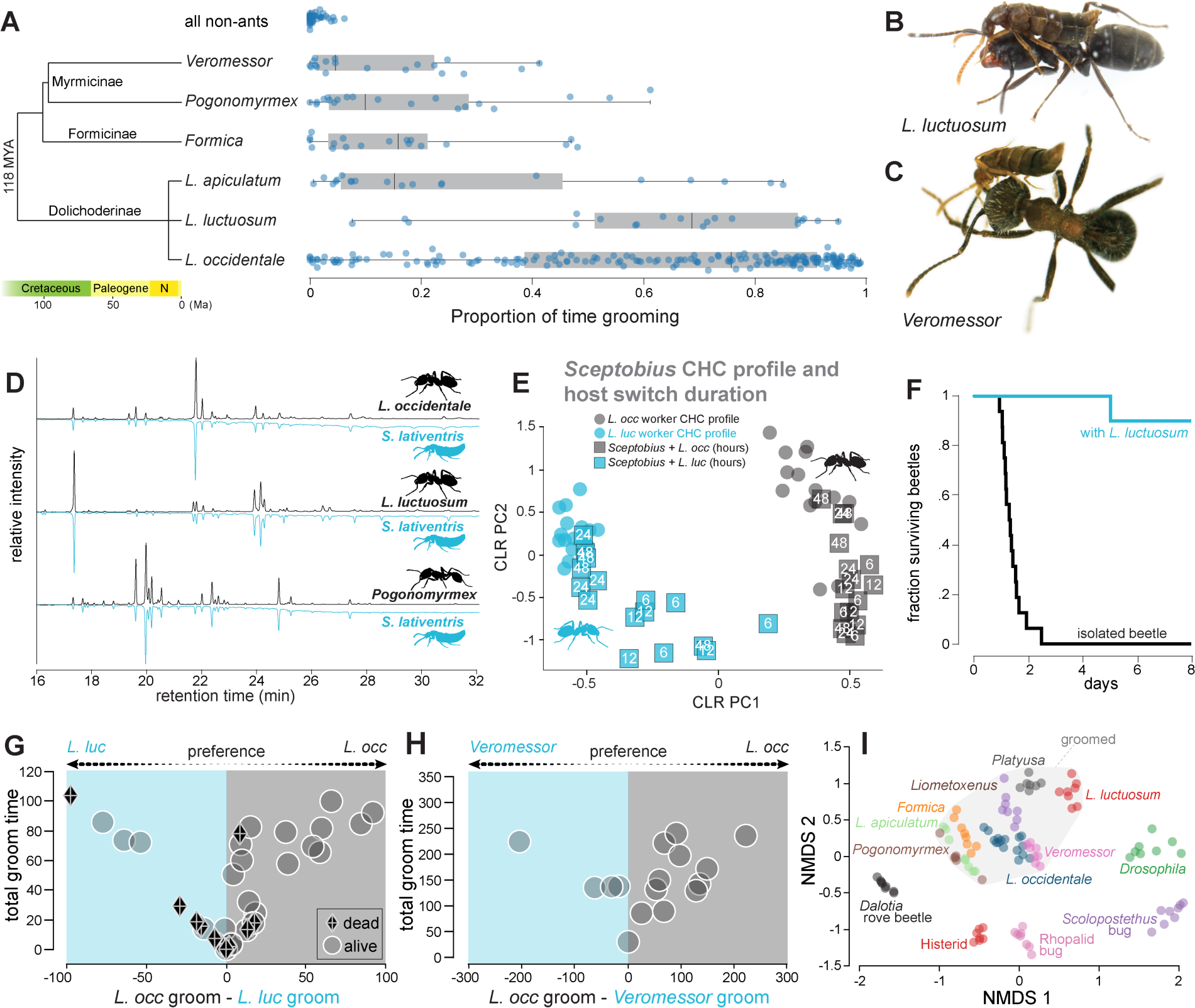
*Sceptobius* shows host promiscuity and negligible chemosensory specialization on its natural host. (**A**) Diverse ant species release *Sceptobius* grooming behavior. (**B, C**) Grooming non-host ants *L. luctuosum* (B) and *Veromessor pergandei* (C). (**D**) Grooming results in turnover of CHCs to match non-host ants. (**E**) After 24 hours of interacting, beetles match the non-host ant’s CHC profile nearly perfectly, and are intermediate in profile between the two ant species after 6–12 hours. (**E**) *Sceptobius* dies rapidly when removed from *L. occidentale* colonies (black line) but can survive inside colonies of *L. luctuosum* once integrated (blue line). (**G, H**) Beetles show a weak preference for host ants in a two-choice assay with non-hosts *L. luctuosum* (**G**) and *Veromessor* (**H**); however, the preference disappears when beetle chooses between dead host and non-host workers (diamonds). (**I**) CHC composition analysis reveals a cluster in chemical space of all the ants and groomed animals, demarcated by the grey convex hull.

*Sceptobius* therefore can and will break its natural partner fidelity when presented with a novel ant species. We asked whether, when faced with a choice, *Sceptobius* displays a preference for its host. We developed a head-to-head preference assay to quantify grooming of host versus non-host ants (**Fig. S5A, B**). In a choice between single workers of *L. occidentale* and *L. luctuosum*, on average, beetles spent slightly more time grooming their host (**Fig. 4G**); however, they still spent substantial time grooming non-hosts, often alternating their grooming between the two ant species. (**Fig. 4G, Fig. S5C, Video S17**). Moreover, this apparent preference disappeared when dead host and non-host ants were provided, suggesting that the response of the non-host ant to attempted grooming, rather than beetle preference, may drive this small difference in groom time (**Fig. 4G**). Remarkably, the beetle still showed only a minor preference for its host over a phylogenetically distant non-host (*Veromessor*), performing long grooming bouts on this ant even when an *L. occidentale* worker was available to groom instead (**Fig. 4H, Fig. S5D, Video S17**). An absolute preference for host over non-host workers therefore cannot explain the natural host specificity of *Sceptobius*. Analysis of the CHC profiles of the ant species groomed by *Sceptobius* revealed an ‘ant cluster’ in chemical space, which also encompassed the two myrmecophile beetles that *Sceptobius* groomed, but excludes all insects that *Sceptobius* ignored (**Fig. 4I**). We conclude that the beetle’s recognition system identifies ants but cannot discriminate hosts from non-hosts in this chemical space. The beetle recognizes non-hosts as potential partners, and its symbiotic behaviors also achieve chemical integration with non-hosts.

We explored whether *Sceptobius* displays specificity for host foraging trails. We allowed field-collected workers of the sister ant species, *L. luctuosum*, to lay foraging trails in an arena before removing the ants. *L. luctuosum* trails are also composed of iridodial and nepetalactol, but in different ratios, and possibly comprise different stereoisomers (**Fig. S6A**). As with trails laid by its host, *S. lativentris* followed naturally laid *L. luctuosum* trails (**Fig. S6B;** movement of free-living *Dalotia* again showed no correspondence with the *L. luctuosum* trail). We also painted crude chemical extracts of the sister ant species in circles and found that *S. lativentris* robustly followed these trails (**Fig. S6C**). We presented the beetle with a choice by painting abutting semi-circles of *L. luctuosum* and *L. occidentale* extracts (**Fig. S7A, B**). *S. lativentris* preferred its host’s trail when it was at higher concentration than extracts from the sister ant species (**Fig. S7C, D**). However, as soon as we switched the sister ant extract to higher concentration, the preference flipped (**Fig. S7C, D**). This result indicates that the trail concentration drives preference, and the beetle simply follows the higher concentration trail regardless of whether it was laid by the host or the host’s sister species. The identities of the iridoids do, however, seem to matter: *Sceptobius* did not follow trails of another dolichoderine ant, *Linepithema humile*, consisting of the iridoids dolichodial and iridomyrmecin^111^ (**Fig. S7E, F**). We infer that *Sceptobius* follows iridodial/nepetalactol trails specifically but cannot distinguish between trails made by different *Liometopum* species.

## Non-hosts and spatial barriers enforce host specificity

Our findings demonstrate that, despite associating with just a single host ant species in nature, *S. lativentris* has a latent promiscuity to associate with a diversity of non-host species. The symbiotic behaviors *Sceptobius* enacts that ordinarily connect it to its host ant are readily performed with non-hosts, and the beetle shows limited preference for its host when given a choice. The cue space that releases symbiotic behavior from *Sceptobius* therefore cannot, by itself, explain the beetle’s extreme partner fidelity. Additional forces must prevent the beetle’s promiscuity from being realized in natural contexts, constraining its association to *L. occidentale* alone. To identify what these forces might be, we created an agent-based model^117^, capturing critical aspects of *Sceptobius* biology that influence its interactions with ants. Using this model, we asked what conditions promote versus repress host switching between nests of different ant species. We then recreated our in-silico model with living insects to experimentally test these findings. Our model is built around three core parameters:

**1. Intrinsic mortality.** *Sceptobius* dies rapidly when isolated from ants, with heightened mortality in dry environments (**Fig. 5A**). A major cause of death is the loss of desiccation-preventing CHCs, which isolated beetles can no longer acquire via ant grooming (**Fig. 5B**). Beetles in humid arenas are partially protected from the hazards of CHC loss, but still die within ∼2 days of isolation. We encoded this information in a parameter, *ΔCHC_loss_* (**Fig. 5D**). Simulated beetles lose CHCs at a specified linear rate when isolated from ants and die when CHCs are depleted. On re-encountering ants, they replenish their CHCs if they survive the interaction (**Fig. 5D**). Wrapped into *ΔCHC_loss_* are other, presently unknown, ant-dependent physiological processes that contribute to the rapid, intrinsic mortality of isolated beetles.
**2. Extrinsic mortality.** Ants are innately hostile to insects with CHC profiles different to their own*. Sceptobius* has an acquired chemical identity and normally possesses an *L. occidentale* CHC profile; consequently, an important observation is that non-host ants exhibit hostility to the beetle. Indeed, to uncover the myrmecophile’s latent promiscuity for non-host ants (e.g. **Fig. 4A**), it was necessary to first obstruct the mandibles of non-host ants in these experiments, thereby enabling *Sceptobius* to groom them. Without doing so, non-hosts killed most beetles within 30 minutes of a grooming trial (**Fig. 5C**). Conversely, *L. luctuosum*, which possesses largely the same CHCs as *L. occidentale*, but in different ratios, is relatively less aggressive to *Sceptobius* than the phylogenetically (and chemically) divergent non-hosts, *Veromessor*, *Pogonomyrmex*, and *Formica* (**Fig. 5C**). We encoded the degree of mismatch between host and non-host CHC profiles in a parameter, *ΔCHC_ID_*, where a greater value increases the likelihood a non-host ant will kill *Sceptobius* in an encounter before the beetle can acquire its CHCs (**Fig. 5E**). The degree of aggression is modelled as a function of the total amount of CHCs (ΣCHC) on the beetle’s body—higher amounts being more detectable, and more likely to elicit aggression (**Fig. 5E**).
**3. Inter-colony distance**. Ant colonies are separated by topographically complex natural terrain. We encoded linear distance between nests in the parameter *Δdistance*. As linear distance between nests increases, the path length that a randomly walking beetle takes to find a new nest in two dimensions increases as a polynomial, and would hence increase especially steeply for topographically complex three-dimensional substrates (**Fig. 5F, G**).

**Figure 5:**
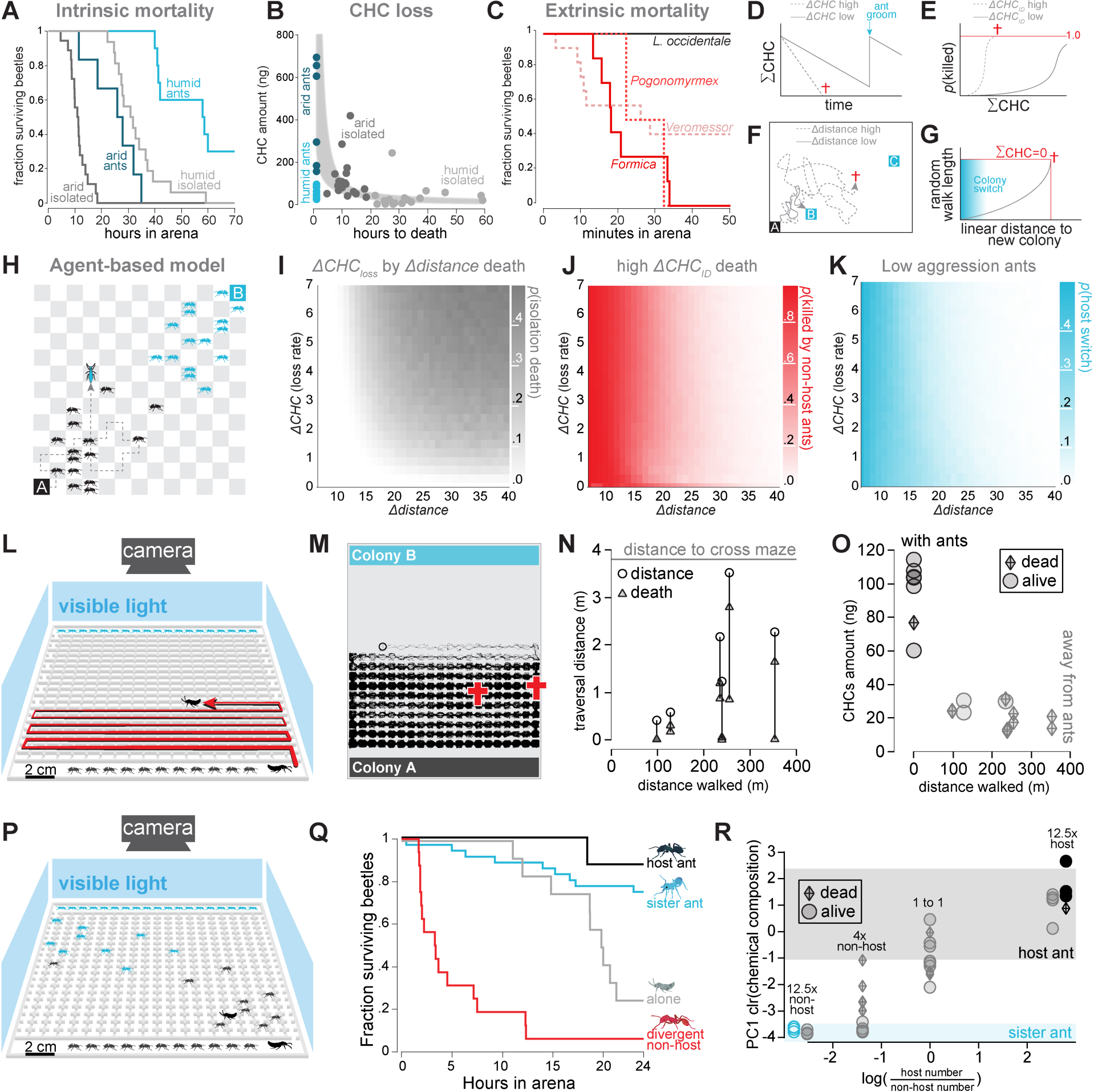
Agent-based host specificity. (**A**) Beetles rapidly die when isolated from host ants. (**B**) CHCs drop steeply on isolation from ants. Grey shaded area shows a 95% CI for a regression of exponential CHC loss calculated via non-parametric bootstrapping. (**C**) Non-host ants quickly kill beetles when their mandibles are unimpeded. (**D-G**) Illustrations of model core parameters. (**D**) Beetles lose CHCs with a loss rate *ΔCHC*, and regain CHCs when they groom an ant. (**E**) Probability of death in an ant encounter scales with *ΔCHC_ID_* as a function of total CHCs on the beetle’s body (ΣCHC). (**F)** As linear distance between ant nests increases (colony A → colony C instead of colony B), the path length for randomly walking beetles increases non-linearly. (**H**) The likelihood that the condition ΣCHC=0 will be met becomes higher with path length. (**H**) Agent-based model: host and non-host ants move randomly through a discretized landscape from opposing colonies; beetles exit the host colony and lose CHCs but can groom and regain them, die from CHC loss or ant aggression, or switch to neighboring ants of varied aggression. (**I**) The model predicts that beetles die alone from desiccation as distance between ant colonies increases. (**J**) Aggressive ants kill beetles, preventing host switching (**K**) Low aggression ants permit host switching when colonies are close together. (**L**) Zig-zag maze arena to test impact of distance on movement to new nest. Beetles can leave host colony and enter opposing colony, but ants are prevented from entering arena. (**M**) Traces of two beetles in maze arena (black) show wandering but no successful arena crosses. The † symbols indicate positions at which beetles died. (**N**) None of twelve beetles successfully crossed the maze (shortest traversal path ∼4 meters), despite wandering hundreds of meters within the arena. (**O**) Beetles in maze arena rapidly lost CHCs while wandering from ants, matching the model. Most died in less than a day away from ants. (**P**) Interaction landscape to test host-switch potential to high- or low-aggression non-host ants. (**Q**) High-aggression ants rapidly kill beetles, whereas low-aggression sister ants do not. (**R**) CHC composition analysis shows that beetles gain non-host chemicals proportional to the ratio of host to non-hosts in the arena, with total integration contingent on the beetle’s survival.

We instantiated a virtual landscape comprising a grid of ‘forest floor’ tiles, with spatially separated colonies of host and non-host ants (**Fig. 5H**). *Sceptobius* beetles dispersing from the host colony interact with host and non-host ants following the rules defined above, and lose CHCs at a rate *ΔCHC_loss_* if unable to encounter and successfully groom an ant. We performed an extensive parameter screen to identify conditions that prevent or favor beetles switching to the non-host colony. The model predicts three regimes: First, by keeping host and non-host ants chemically similar (*ΔCHC_ID_* = 0.1) but modulating landscape area, we find that beetles fail to host switch in large landscapes, and instead die from intrinsic mortality (*ΔCHC_loss_*) when ant nests are far apart (**Fig. 5I**). Conversely, when inter-colony distances are short, high aggression from chemically divergent ants prevents host switching (**Fig. 5J**). Finally, in scenarios where CHC divergence of non-hosts is relatively small, beetles can successfully host switch, contingent on non-host nests being spatially close enough to avert the beetle’s intrinsic mortality (**Fig. 5K**). These outcomes indicate that the natural host specificity of *Sceptobius* could in theory hinge on forces that are independent of the beetle’s agency. Rather, its strict association with *L. occidentale* may emerge from coarse-grained attraction to general ant CHCs combined with external enforcement, firstly from dispersal constraints imposed by the environment outside of colonies (which promote the beetle’s intrinsic mortality away from ants), and secondly from deleterious behavioral interactions with non-host ant species (should these interactions arise). The model also underscores how host specificity is probabilistic rather than absolute, with certain circumstances elevating the chances that barriers will break down and switching will result.

We attempted to empirically replicate these findings by constructing large-scale, host-switching arenas in which we could track beetle behavior across a landscape with colonies of host and non-host ants (**Fig. S8A, B**). We first tested the prediction that beetles would die when isolated from ants when attempting to cross a navigation-cue-free-space to a new ant nest. We introduced a zig-zag course and tracked whether beetles crossed from one *L. occidentale* colony to another, while selectively preventing the ants themselves from entering the arena (**Fig. 5L, Fig S8A**). Even though the minimal path length to cross this maze was only ∼4 meters, no beetle successfully crossed the maze, despite typically wandering for well over 100 meters (**Fig. 5M, N**). Net movement of most beetles was <2 meters along the course, and the majority ended up dying less than a meter from their starting colony (**Fig. 5N**). Total CHC levels on the bodies of these dispersing beetles decreased massively after leaving their parent colony (**Fig. 5O**), consistent with the assumption of our model. Together, these data confirm that even a small linear distance between ant nests creates a near-insurmountable physical barrier for the beetles to navigate, precluding them from switching between nests in the absence of navigational cues. Moreover, as linear distance between nests increases, the area of a topographically complex space the beetle must explore balloons, yielding a nearly infinite walking distance. On exiting a colony of its natural host, *L. occidentale*, however, navigational cues exist in the form of iridoid foraging trails, permitting long distance movement but acting to restrict *S. lativentris* to nests of this ant alone.

We next tested the prediction that, when in close enough spatial proximity to feasibly host switch, ants with strongly dissimilar CHCs (high *ΔCHC_ID_*) would kill beetles, thereby aggressively rather than spatially enforcing specificity. To do this, we built a second arena with a simplified spatial structure through which both beetles and ants could move freely and interact, emerging from their source colonies located at opposite ends of the arena (**Fig. 5P, Fig S8B**). We performed trials with different starting numbers of host and non-host ants of different species. Within hours, phylogenetically distant non-host ants with strongly divergent CHCs ants killed all beetles (**Fig 5Q**). We observed similar results with three other species of CHC-divergent ants, confirming that even if the beetle reaches a non-host ant colony, its previously acquired *L. occidentale* CHC profile is likely to trigger aggression and prevent host switching.

Finally, we examined whether a low *ΔCHC_ID_* with a potential new host might allow *Sceptobius* to host switch. Employing the chemically similar congeneric ant, *L. luctuosum*, we observed high survival rates for *S. lativentris* when permitted to interact with this sister ant species across a range of host: non-host ant ratios (**Fig. 5Q**). Strikingly, even in the case of 20 host and 250 non-host ants—in which all host ants were killed by the sister ant species—all the beetles survived, groomed the non-hosts, and switched to the *L. luctuosum* colony after acquiring the non-host’s CHC profile (**Fig. 5R**). Strikingly, we found the higher the ratio of non-hosts to hosts, the closer the CHC profiles of the beetles to the non-host became (**Fig. 5R**). Beetles with intermediate pheromone profiles were animals that died during the run, having failed to host switch (**Fig. 5R**). These findings demonstrate that host switching may be possible were *Sceptobius* to encounter a weakly aggressive ant species in close proximity. In the San Bernardino mountains, a case of sympatry of *L. occidentale* and *L. luctuosum* was recently documented^118^. We have since visited this locality and collected numerous *S. lativentris* from multiple *L. occidentale* colonies, but none from *L. luctuosum* nests despite their proximity within tens of meters. We infer that the enforcement barriers identified herein have so far repressed host switching, but predict such a scenario may be plausible in the sympatric range of these ant species. We could not experimentally test one further host switching scenario derived from our model: that if the limit of ant detection of CHCs is higher than the minimum amount needed for *Sceptobius* to survive, then switching to chemically divergent ant species may be possible, though infrequent (**Fig. S8C**). Such a situation implies that beetles with strongly depleted CHC levels may be able to overcome enforcement barriers—their survival rescued by chance host switching to a diversity of potential ant species.

## Discussion

Knowledge of the sensory information that connects symbiotic organisms to their hosts is fragmentary; so too is an understanding of the forces that shape the often-strict fidelity of these partnerships. Using a tractable ant-myrmecophile model, we have identified ant-derived cues that are exploited for host recognition and long-range dispersal by the myrmecophile. Surprisingly, we uncovered a pronounced lack of chemosensory preference of the myrmecophile for its host, manifested in its near-equivalent ability to use corresponding sets of chemical cues from alternative ant species. Hence, despite these ant compounds possessing many species-specific features^104,105^, they do not underlie the observed, stringent specificity of the myrmecophile towards its single ant host. Instead, we found that rapid mortality coupled with an inability to disperse to new ant nests without long-range dispersal cues strongly spatially enforce the symbiont’s host association (**Fig. 6**). Additionally, hostility of alternative ant species towards *Sceptobius* when coated in its natural host’s CHC profile limits its realized host range (**Fig. 6**). We demonstrated through simulation and subsequent experimental testing that these barriers suffice to make host switching rare, and can enforce the association of the beetle to a single ant species. We cannot rule out that presently unknown host-derived cues may exist that attract *S. lativentris* to its natural host over alternative ant species (e.g. from nest material^119^, so far not studied by us). Nor can we be certain that *S. lativentris*’ life history is fully compatible with alternative ants (though we hypothesize compatibility with at least congeneric ants that are biologically highly similar to its natural host). Regardless, our findings show that even if such impediments to host switching exist, the enforcement mechanisms we identify are themselves a major initial barrier, capable of restricting the symbiont’s range to a single ant species despite its lack of host preference.

**Figure 6:**
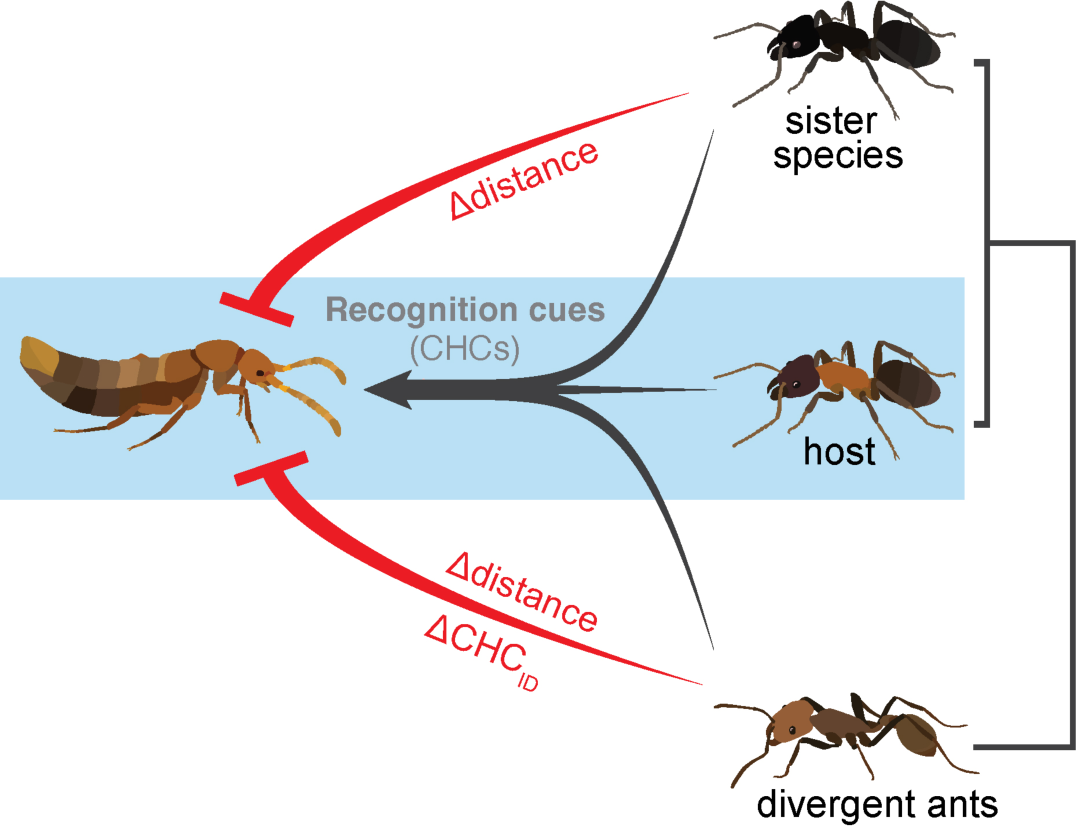
Forces shaping myrmecophile host specificity. *Sceptobius* possesses a coarse-grained ability to detect ant CHCs and enact grooming behavior, conferring a latent promiscuity to switch to other ant species. However, insurmountable spatial barriers prevent switching to phylogenetically close, chemically similar ant species, and both spatial barriers preclude switching to phylogenetically divergent, chemically dissimilar ants.

Enforced specificity contrasts with models that invoke sensory tuning to host-derived cues. We note that these models have emerged primarily from studies of vagile specialists (e.g. flight-capable plant- or blood-feeding insects). For such organisms, an abundance of competing environmental stimuli may necessitate sensory tuning, limiting interactions with off-target hosts. Conversely, we propose that enforced specificity may be a key determinant of host ranges for intimate symbioses, such as host-embedded forms of parasitism^6,52^. In these systems, the potential for interactions with alternative hosts is low, imposing weak selection for partner discrimination. Even for more mobile specialists, however, external enforcement may still play a critical role. In the case of phytophagy, for example, toxins from plant secondary chemistry^120^, and inadequate defense against natural enemies^47^, may exert analogous restrictions on diet breadth and hence function as an early and sustaining force behind the evolution of sensory tuning. In effect, *Sceptobius* represents the counterpart to these systems—a natural experiment that reveals what happens when specialization evolves in the relative absence of alternative hosts. It follows that some of the most tightly integrated symbionts may be those most prone to experiment with alternative hosts, should they encounter them.

The beetle’s latent attraction to novel hosts can be viewed as a non-adaptive trait that is often deleterious when realized, leading to beetle death, and potentially neutral with regards to fitness should the beetle successfully host switch. Ecological fitting between symbiont and novel host will dictate whether the new partnership attains evolutionary stability^121,122^. Should these encounters arise sufficiently frequently, we predict host switching will ultimately occur. Highly specialized symbionts, including endo- and ectoparasites, many parasitoids, and social parasites like myrmecophiles, are profoundly host-dependent. Obligate entrenchment in the biology of another species places these organisms at high risk of co-extinction^123,124^. Nevertheless, many ancient radiations of such symbionts exist. Almost invariably, patterns of host use across their phylogenies reveal a historical record of host switching (e.g. ^87,96,98,125–130^), and evidence obtained from some groups implies that current host ranges may, in part, emerge under enforcement by repressive actions of alternative hosts^65,66,84,131–133^. We suggest that ancient, obligately symbiotic taxa are the outcome of macroevolutionary survivorship bias for clades with intrinsic promiscuity: this property potentiates host switching during chance events when enforcing constraints have been overcome. Latent promiscuity, though possibly non-adaptive, may be crucial to the deep-time persistence of symbiotic lineages.

## Acknowledgements

We thank James Danoff-Burg for early and invaluable insights into the biology of *Sceptobius*, and Michael Dickinson for guidance with initial behavioral arena designs. We are grateful to Christiane Weirauch (UC Riverside) for assistance with hemipteran identification, John Truong for ant husbandry, and members of the Parker laboratory for assistance with fieldwork. This study was supported by an Army Research Office MURI grant to JP and JB (W911NF18S0003), with further funding to JP from NSF (CAREER 2047472), NIH BRAIN initiative (NINDS R34NS11847), an Alfred P. Sloan Fellowship, Pew Biomedical Scholarship, Rita Allen Scholarship, Klingenstein-Simons Fellowship, and research Grants from the Shurl and Kay Curci Foundation and Okawa Foundation. JMW is an NSF GRFP recipient and also received funding from a Graduate Research Excellence Grant (Rosemary Grant Advanced Award) from the Society for the Study of Evolution.

## Data and code

All data and code (in Jupyter notebooks) used to generate the figures in this paper can be found online at CaltechData: https://data.caltech.edu/records/6z3fs-sm018

## Methods

### Specimen collection and husbandry of *S. lativentris* and *L. occidentale*

Beetles and ants were collected in the Angeles National Forest, primarily near the parking lot of Chaney trail and along Millard creek in Altadena, CA (34.2163413, -118.146500), and near Gould Mesa Trail camp, along Gabrieleno trail, also near to a creek (34.2222252, -118.1785464). *Liometopum occidentale* builds nests in the bases of oak (*Quercus*, especially *Quercus agrifolia* at the listed collecting sites) and bay trees (*Umbellularia californica*). In warmer/dryer conditions (especially during the summer) leaf litter near to ant nests and along foraging trails was sifted, and the trays examined for beetles. During colder weather and early in the spring, beetles walk on the trees housing the ant nest, often near the nest opening. Blowing exhaled air into an undisturbed nest often increases activity, enabling collection of *S. lativentris* exiting the nest. Beetles were captured via aspirator and placed with host ants in falcon tubes with slightly dampened KimWipes. *S. lativentris* are abundant, and could be collected during most of the year, but are more difficult to find during November–January. Collecting expeditions yielded as few as zero beetles on the coldest days, compared with up to ∼200 beetles per colony per day during later spring and summer. To keep *S. lativentris* in the laboratory, beetles were housed with an excess of well-fed *L. occidentale* workers collected from the same colony that yielded the beetles. In the laboratory, beetles were placed with host ants into ∼10 inch x 10 inch Rubbermaid boxes with a Fluon barrier (2/3 water, 1/3 Insect-a-Slip) painted on the sides to avoid escape. Animals were provided a feeder of hummingbird nectar (4 parts water, 1 part nectar) and a test tube setup with water and cotton balls to provide moisture. Specimens housed this way survived up to several months in the laboratory. *Sceptobius lativentris* were sexed for experiments based on a dimorphism in antennal setation: males have a high density of spatulate setae on antennomere 3, whereas females have few such setae. Ice was used as an anesthetic for sexing beetles under the microscope and sorting them for experiments (CO_2_ anesthesia was found to cause mortality).

### Grooming behavior arenas

**Grooming Arena 1:** A multiplexed array of circular arenas was constructed to record grooming behavior (**Fig. S1A**). To avoid vision influencing behavior, behavioral arenas were constructed out of 1/8^th^ inch infrared-transmitting acrylic (Plexiglass IR acrylic 3143) which transmits far red and infrared while blocking visible light. Hence, experimental trials were conducted in darkness. Arenas consisted of a base layer of finely wet-sanded acrylic (to provide a texture on which beetles could walk), on top of which was placed a second layer with multiple, 2 cm diameter, circular wells. Finally, a top acrylic roof layer was added to contain the animals inside the arena. Slight modifications of these 2 cm arenas were used throughout the data collection period, with either fixed-well shape or a sliding door design to allow a particular start time for insect interactions. Behavioral interactions were run in a dark incubator, situated within a dedicated behavior room with the lights switched off, behind a blackout curtain to further ensure that the insects were behaving in complete darkness. Arenas were backlit with a custom-built IR850nm LED PCB and diffused with a semi-opaque white acrylic sheet. Recordings of interactions were made using a Flir machine vision camera (BFS-U3-51S5M-C: 5.0 MP) at 3 frames per second with a Pentax 12mm 1:1.2 TV lens (by Ricoh, FL-HC1212B-VG), for 6 hours.

**Grooming Arena 2**: Later, similar arenas as above were also built (**Fig. S1B)**, but designed with side rather than top IR illumination, a higher camera frame rate, and higher resolution per experimental well to better maintain visibility of the beetle when grooming during trials, and provide more information-rich behavioral data (this higher spatial-temporal resolution data was unnecessary for the present study, but was gathered with futures studies in mind). For this second setup, an 8-well arena with similar design as mentioned above was used, with a base layer of sanded IR acrylic, a wall layer with eight 2 cm circular arena cutouts, a ceiling of static dissipating acrylic with a rim of IR acrylic, and a second roof of IR-transmitting acrylic. An aluminum frame to hold the arena was constructed with 1” T-slotted framing from McMaster-Carr, along with open gusset structural brackets, and custom laser-cut ¼” acrylic brackets. IR flood lights (Univivi U6R) were side-mounted, and a Flir machine vision camera (BFS-U3-51S5M-C: 5.0 MP) with a Pentax 12mm 1:1.2 TV lens (by Ricoh, FL-HC1212B-VG), was used to record at 60 frames per second. An Arduino-based external trigger was also used to maintain the frame rate of the camera. The arena was placed in a dark, temperature-controlled incubator set to 18 °C. A thermal camera (Flir Lepton 2.5 with Flir Purethermal-2), was used to determine that the arena itself, heated by the IR lights, maintained a consistent temperature of 21 °C during the trials.

**Loading arenas and preparing behavioral experiments:** To control for sex differences, only male *S. lativentris* were used for all grooming experiments (although female beetles exhibit overtly identical grooming behavior). Beetles were isolated in a container with two moistened KimWipes for 30 minutes–1 hour before loading into behavioral arenas. Beetles and interactor ants/insects were anesthetized on ice for 10 minutes before loading into a pre-chilled arena in a 4°C refrigerator to prevent them escaping. Loaded arenas were then placed into the incubator setup as described above and recording started. In the case of moving/sliding door arenas, arena pieces were slid together to start interactions after *S. lativertis* started moving around its arena well, ∼10 minutes after beginning the loading process.

### Machine learning analysis of grooming behavior

**DeepLabCut for grooming arena analysis:** DeepLabCut^107^ was used to track beetle and ant/other insect behavior. A network model with five labeled points on the *S. lativentris* and five labeled points on each interactor was used (**Fig. S1C**). We found that a single model to detect these key points could be trained to identify the beetle and the other insect, regardless of the species by including additional training frames to the dataset for each interactor type. A ResNet50 network architecture was trained and subsequently used for annotation. The final network was trained on ∼2300 frames from more than 200 videos. This network achieved an error of 2.53 pixels for the training data, and 4.45 for the test data, which represents an error of less than 1/5^th^ of a mm within the arena (less for most videos). If no detection for a given animal was present in a frame, linear interpolation from the last known position to the next known detection position was used to fill the gap. The distance between the beetle and the other interactor insect was calculated during the trial, and an interaction was considered a grooming bout if the beetle was within 3 mm of the ant for at least 30 seconds.

**YOLOv8 preference assay analysis:** To test whether *S. lativentris* showed a preference for grooming its host ant over other ants, a single *L. occidentale* host ant worker and either a single sister ant (*L. luctuosum*) or a phylogenetically divergent ant (*V. andrei*) were placed with a single beetle in an arena well. To quantify the relative preference for one ant species over the other, the amount of time the beetle spent grooming each ant during a two-or six-hour trial was determined. For analysis, behavioral videos were thinned to one frame per 16.7 seconds. YOLOv8^134^ was used for detection and bounding box generation of the location of each ant and each beetle during the behavioral trial (**Fig. S5A, B**). Frames were extracted uniformly from each behavioral trial video (10 per video for the *L. luctuosum* analysis for a total of 480 frames from 48 trials, or 30/31 per video for the *V. andrei* analysis for a total of 481 frames labeled). An 85% training data -15% validation data split was used. Labeled data with a bounding box per animal were manually generated in CVAT (https://www.cvat.ai/). The network was trained with YOLOv8’s default settings (epochs: 100, patience: 50, batch: 16, imgsz: 640, lr0: 0.01, lrf: 0.01, momentum: 0.937, weight_decay: 0.0005, warmup_epochs: 3.0, warmup_momentum: 0.8, warmup_bias_lr: 0.1, etc.) (see **Fig. S5A**, **B** for training results). Detection was then performed on all frames of the thinned behavioral videos. For each frame, the highest confidence detection for each animal type per frame was taken. If no detection for a given animal was present in a frame, linear interpolation from the last known position to the next known detection position was used to fill in gaps. The distance between the beetle and the other interactors during the trial was calculated, and considered a grooming interaction if the beetle was within 3 mm of the ant for at least 90 seconds. To estimate the amount of time grooming each individual ant type, ambiguous grooming bouts where the beetle was within 3 mm of both ants were eliminated, and the cumulative time spent grooming just one or the other ant species unambiguously was calculated. To quantify preference, total groom times for each ant species were subtracted to obtain a differential groom time estimate.

### Trail following analysis

**Naturally laid trail arena:** To assay trail-following ability and specificity of *S. lativentris*, a large (16 × 20 inch) open field behavioral arena was constructed, enclosed within IR-transmitting acrylic (**Fig. S3A**). To provide a naturalistic ant-trail stimulus, ants from a laboratory colony of *L. occidentale* were allowed to lay down a trail in the arena, with a large sheet of filter paper covering the bottom of the arena and acting as a diffuser for the IR 850nm strip backlights. After starving the ants for 2 days, the colony was connected to the arena environment, with a foraging object (sugar water) available at a distal region of the arena (**Fig. S3A**). Within the free field arena, obstacles were placed to force the ants to lay a trail with a specific geometry. After allowing the ants to forage for 12 hr, the ant colony was disconnected, the arena was filled with CO2 to anesthetize remaining ants, and all ants and obstacles were removed from the arena (**Fig. S3C**). The trail-bearing filter paper was placed back into the arena, and single *L. occidentale* workers (**Fig. S3D**), *S. lativentris* beetles (**Fig. S3E**), or free-living *Dalotia coriara* beetles (**Fig. S3F**) were then placed into the arena. Insect movement traces were recorded with a Flir machine vision camera (BFS-U3-51S5M-C: 5.0 MP) with a Pentax 12mm 1:1.2 TV lens (Ricoh, FL-HC1212B-VG). To quantify trail following of individual insects, net frame-to-frame movement in the arena was correlated with ant movement flow at that position in the arena. Frame-to-frame beetle or ant movement was calculated based on thresholding the difference between subsequent frames to find locations of flow. In addition to quantifying net movement of the beetles via frame-to-frame difference, blob tracking on beetle position throughout a behavioral trial was also performed. For this, median filtering was carried out on a set of frames from the beetle-walking-in-trail-arena video to construct a background frame. With OpenCV, blob detection was performed on background-subtracted frames from the video. The median position of the blob was used to make a trajectory for beetle position in the arena.

**Multi-well trail arena:** To probe the chemicals relevant for trail following, a multiplexed assay to test beetle behavior in response to artificially applied trails was also developed (**Fig. S4A**). An arena with nine square wells of 3.5 inches × 3.5 inches was constructed. The arena was constructed from stacked layers of acrylic. The base was ¼ inch clear acrylic, with an a 1/8^th^ inch thick layer of Plexiglass IR acrylic 3143 placed on top to block visible light. On to this was placed an IR-transmitting acrylic layer with a 12-inch × 12-inch opening that fitted a 12-inch × 12-inch square of 1/8^th^ inch thick glass with a ground surface to provide grip for beetles to walk. An opaque white acrylic layer with nine wells of 3.5 by 3.5 inches was placed onto this, with fluon applied to the walls of each well to prevent insects from climbing onto the roof. The roof was placed over this, comprising a 1/8^th^ inch layer of static-dissipating acrylic with a further 1/8^th^ inch layer of IR acrylic to block visible light, and an additional 1/4^th^ inch layer of clear acrylic to weigh down the ceiling and keep it flat. The layers were all held together by screws affixed to a metal frame and backlit with IR 850nm strip LED lights. The arena was monitored with a FLIR machine vision camera (BFS-U3-16S2M-CS: 1.6 MP) with a Pentax 12mm 1:1.2 TV lens (by Ricoh, FL-HC1212B-VG). Extracted ant compounds were painted onto a ground glass floor in a circular pattern within each well. Behavioral trials within this arena were two hours long.

To analyze the resulting videos, OpenCV was used. First, videos were cropped to extract individual wells from the array. Individual wells were warped to square them from any small camera distortions and set to a constant resolution of 320 × 320 pixels per well. A background frame was then constructed via median filtering of a set of images from the given well. Background subtraction was then performed for each well, and the OpenCV blob detection method was used to threshold each frame and locate the beetle (**Fig. S4B**). The position of the beetle in the well was saved. To calculate the degree of trail following observed in the trial, the positional information given by blob tracking was used, and circular arcs within the animal trajectory extracted. To do this, regions of interest were defined as sections along the circular chemical trail of approximately 0.5 cm, moving along the circle and diverging from the trail by ∼0.5 cm either side of the path of the circle. Twelve such regions were defined per circle, at intervals of 30° along the circle (**Fig. S4B**). Instances where an animal traversed through these twelve regions sequentially, from one to the next, for an entire revolution of the circle were measured. Each such traversal was counted as a single circular trail following event. The distance traveled while the animals were traversing these circles was then calculated.

**Preference assay in trail arena:** To test whether *S. lativentris* prefers trails of its host ant over its sister ant species, a variant of the multi-well trail arena was used (**Fig. S7A**). An approximate concentration match of crude extract from the host ant *L. occidentale* or the sister species *L. luctuosum* were prepared. To do this, hundreds of ants of the two species were extracted of the two species in hexane, and an aliquot of 2 microliters injected into a GCMS; the region of a GCMS trace (GCMS methods described elsewhere) representing the iridoid fraction of the trace was integrated to approximate the concentration of these compounds. These values were used to mix approximately equal concentration solutions of each extract. A dilution to 1/5^th^ the concentration was also made to generate a comparably low and high concentration extract for the host and sister ants. Abutting lobes of semi-circular trail were then painted with low or high concentration of extract from the two different ant species. A single beetle was then placed in each arena with the high-low concentration trails and its movement was recorded for a two-hour trial. For this assay the same setup as described above for the multi-well trail arena was used (**Fig. S4A**), but with a BFS-U3-63S4M-C 6.3 MP camera with a Pentax C61232KP 12mm F1.4 Manual Lens with Lock Screw. To quantify the results, movement of the animals during the trial, the cumulative pixel difference between subsequent frames was calculated for the whole experiment. Regions of interest (ROI) were then manually defined as the arms of the trail lobes belonging to either species. The total movement in each of these ROIs was summed and subtracted to see the difference in total movement during the trial on one trail lobe or the other, which was used as a trail preference index. Example depictions of the total movement histogram and the movement histogram for a given trail lobe ROI are shown (**Fig. S7B**).

### Identification of chemical compounds and analysis of CHC profiles

For identification and profiling of chemical compounds from insect species used in this study, we used gas chromatography mass spectrometry methods described previously in Brückner et al (2021)^114^ and Kitchen et al (2024)^113^. Specimens were freeze killed at -80°C and stored at -20°C until extraction. Compounds were extracted by submerging individual insects in 70 microliters of hexane (SupraSolv® n-Hexane, Merck) containing 10 ng/microliter octadecane (Sigma Aldrich) as an internal standard. After 20 minutes, the hexane was transferred to a 250µL glass small volume insert and samples were either analyzed immediately or stored at -80°C until analysis. Analysis was performed on a GCMS-QP2020 gas chromatography/mass-spectrometry system (Shimadzu, Kyōto, Japan) equipped with a Phenomenex (Torrance, CA, USA) ZB-5MS fused silica capillary column (30 m x 0.25 mmID, df=0.25 µm). Samples were injected (2µL) into a split/splitless-injection port operated in splitless-mode at 310°C. Helium was used as a carrier gas with a constant flow rate of 2.15 mL/min. The column was held at 40°C for 1 min, ramped at 20°C/min up to 250°C, ramped at 5°C/min up to 320°C, and then held at 320°C for 7.5 minutes. The transfer line and MS ion source temperatures were kept at 320°C and 230°C respectively. Electron impact ionization was carried out at an ion source voltage of 70 eV, collecting 2 scans/sec from *m/z* 40 to 650.

CHCs were identified based on their fragmentation patterns and retention indices compared to a standard series of n-alkanes^135^, by comparison to library spectra (NIST 14 library), and comparison of retention indices and fragmentation patterns to known compounds in *Drosophila melanogaster*^136^ and *Liometopum occidentale*^108^. Unless specified, the double-bond positions for most alkenes, dienes, and trienes were not determined. Semi-quantification of CHC amounts was carried out by calculating the ratio of each hydrocarbon peak to the C18 internal standard. Absolute amounts and percent composition were calculated for CHCs in all GC traces. Percent composition data were center log-ratio transformed following zero replacement before performing PCA for **Figures 4E** and **5R**. A subset of the total CHCs were used in the PCA analysis. Pairwise Bray-Curtis dissimilarity was calculated for untransformed percent composition data for all samples prior to performing non-metric multidimension scaling (NMDS) for **Figures 2H** and **4I**. NMDS was performed using the metaMDS function from the R package vegan^137^.

Iridoids were identified by comparison to library spectra (NIST 14 library) and by comparison of fragmentation patterns and retention indices to previously described iridoids in ants^138^. Both iridodial and nepetalactol possess multiple stereocenters and while we were able to determine that multiple stereoisomers for both compounds were present, we could not identify the exact configuration of the compounds.

### Fractionation of ant chemical compounds

A bulk hexane extraction of tens of thousands of *L. occidentale* workers was made, with a concentration estimated of ∼50-100 ants per ml. This stock extract was stored at -20°C. 50 ml of extract was concentrated to dryness by rotary evaporation, and the residue re-dissolved in 5 ml hexane. A vacuum flash chromatography column was prepared from a 10 ml sintered glass funnel filled with 230-400 mesh flash chromatography grade silica gel. The silica gel bed was packed with hexane by pulling the solvent through with vacuum. The hexane solution of concentrated ant extract was loaded onto the column, rinsing with hexane. The column was eluted sequentially with:

**a.** 3 x 12 ml hexane
**b.** 3 x 12 ml 5% cyclohexene in hexane
**c.** 2 x 12 ml ether
**d.** 2 x 12 ml EtOAc

This method accomplished the fractionation, leaving saturated hydrocarbons in fraction 1 and 2, unsaturated hydrocarbons in fractions 5 and 6, and polar compounds in fractions 7 and 8. Fractions 1 and 2 were combined, as well as 5 and 6, and 7-10, and volumes adjusted to 10 ml, or ∼250 ant equivalents/ml. The polar and non-polar fractions were used for experiments.

### Agent-based modeling

An *in-silico* agent-based simulation was designed to reproduce *Sceptobius* interactions with host and non-host ants across a virtual landscape. The model incorporates two core limitations on beetle survival: i) intrinsic mortality due to CHC loss and other isolation-related sources of death; and ii) extrinsic mortality caused by encounters with ants that diverge from the beetle’s own, current CHC profile, with greater divergence inversely related to beetle survival probability. We built models initially in Matlab and subsequently in Python, where N × N grids of variable size are instantiated representing a 2D forest floor through which beetles and ants navigate. Host colony (Colony A) and non-host (Colony B) are located at opposite corners of the forest grid. In each simulation, all beetles start within colony A, and all ants start within their respective colonies. The beetles also start with a full supply of CHCs: for the purpose of the model, we encoded two CHCs, one for recognition, and one for resistance to desiccation. Both CHCs are lost at a rate *ΔCHC_loss_* when beetles are isolated from ants, the value of which was varied across in silico runs. At each step of the simulation, each ant moves to one of the four squares contiguous with its current squares. The probability of movement was biased such that ants further away from their source colony were relatively more likely to move backwards to a square closer to the nest. When an ant from Colony A encounters an ant from Colony B, the winning species is determined by a coin flip. If ants from one Colony outnumber ants from the other within a square, the outnumbered ants die. At each time step, the beetles also move within the arena and lose their CHCs at the rate *ΔCHC_loss_*. When beetles encounter an ant with the same CHCs it possesses (*ΔCHC_ID_* = 0), it grooms, re-gaining full quantities of both recognition and desiccation-resistance CHCs. When beetles encounter a novel, non-host ant with divergent CHCs to its own (*ΔCHC_ID_* ≠ 0), its probability of being killed is function of both the degree of CHC divergence (*ΔCHC_ID_*) and the total amount of CHCs on its body (ΣCHC). If the beetle survives the encounter, it replenishes its CHCs and changes its recognition CHC to the identity of the novel, chemically divergent ant. The probability that the beetle survives the encounter is given by:

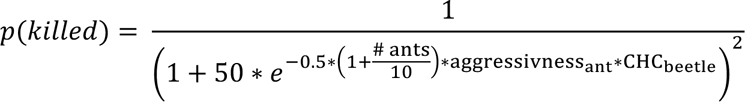

When ants or beetles die, they are reborn at their starting colony to keep the number of animals in the simulation constant. We ran each simulation for 1000 steps, and screened through parameter space, repeating each set of parameter values 100 times to converge on average outcomes for each set of conditions. The simulation was run across a spectrum of CHC loss rates (*ΔCHC_loss_*), degrees of non-host ant aggressiveness (*ΔCHC_ID_*), and different inter-colony distances by varying the forest floor arena area. In each case, we recorded how these parameters influence i) the frequency and cause of beetle death (intrinsic versus extrinsic mortality); ii) the probability beetles successfully obtain CHCs from non-host ants, and iii) the probability beetles were able to host switch to the non-host colony.

### Host-switching behavioral platforms

To experimentally test the in-silico model of host switching, arenas were constructed that matched the design of the model

**Cross arena:** To recreate beetle interactions with opposing colonies of host and non-host ants, we constructed an arena area comprised two nest chambers flanking a central dispersal arena (**Fig. S8B**). Beetles and varying numbers of ants of two different species were placed into the nest chambers and were permitted to disperse into a central area comprising a 20 × 20 grid of cross-shaped separators that formed an array of connected wells in which the ants and beetles could interact. The cross-arena plate design was printed with a Prusa I3 MK2 3d printer in clear PLA. The base of the piece was 1/8^th^ inch thick, and the wall component also 1/8^th^ inch thick. Acrylic was used to sandwich the 3D printed component and provide a ceiling to contain the animals. Screw holes were cut into a base plate of 1/8^th^ inch clear acrylic, matching holes in two 1/8^th^ inch pieces with cutouts of the same dimensions as the arena, and a ceiling of clear acrylic. This created a 4-layer sandwich encasing the arena. Two such arenas were mounted next to each other in a metal frame. We maintained color information in these trials to help differentiate ants of different species and the beetles. We placed white LED photography lights around the arena on four sides and mounted a color camera (BFS-U3-200S6C-C: 20 MP, 18 FPS, Sony IMX183, Color) to the frame with a 16mm 10MP Telephoto Lens for a Raspberry Pi HQ Camera. For experiments, behavioral trials were run for 24 hours at 5 frames per second. When beetles survived or were physically intact enough, their CHCs were extracted, along with two of each ant type from the run, in hexane including a C18 standard for 20 minutes followed by GCMS analysis of the extracts.

**Cross-maze arena:** To test whether beetles could survive/navigate to a new nest of ants without any dispersal cue, a variant of the above arena was constructed in a maze configuration (**Fig. S8A**). Only the end walls connecting the flanking colony chambers to the arena were left open, forcing the beetles to traverse a distance of ∼4 meters at minimum to find a group of ants at the other end of the arena. The ants were retained behind a size-selecting door that would allow the beetle to enter, but which was too small for the ants themselves to pass through. Experiments with beetles in this arena were run for 24 hours, after which the beetles removed and their CHCs were subsequently extracted for GCMS analysis (as above for the cross arena).

**Blob tracking for cross-maze distance analysis:** To analyze the behavioral trials in these arenas, a combination of manual annotation and machine vision methods were used. To calculate the distance the beetles moved in search of ants in the maze, a blob tracking approach was used. Using the python implementation of OpenCV, each individual replicate (right or left arena) was first cropped and de-distorted with the warp perspective method to square the image and correct for the fish-eye effect from the wide angle lens. Frames were downscaled to 20% of their original resolution to speed the blob tracking analysis, giving final dimensions of ∼500 × 500 pixels per arena. From the resulting videos, a background frame was constructed using a median filter on ∼10 frames taken uniformly at times during the first ∼5 hours of the video. After making the background frame, downscaled videos were looped through, background-subtracted by frame, and blobs in the frame detected, with results saved. With the outputs of the blob tracker, detections were run through SORT to generate IDs for the tracked blobs. Short, spurious trajectories where the blob tracker made non-beetle detections were also filtered out with this information. The distances each beetle traveled in the experiment during the run was summed. Trajectories were also used to locate the farthest point in the maze that the beetle reached during the trial. The location where the beetle ended the run was manually annotated. Together, these data provided the total distance traveled, how far beetles penetrated the maze, and the beetle’s position at the end of the trial. The distance traveled was correlated with the CHC level from a 20-minute extraction in hexane (with C18 standard) of the beetle at the end of the trial.

**Manual curation of beetle death times:** For the cross arena and cross-maze arena, time to death for beetles in the experiments was manually annotated. Videos were scrubbed through to locate the last time that the beetle moved in the arena under its own volition (thereby avoiding instances when ants moved dead beetles).

## Supplemental Figure Legends

**Figure S1.**
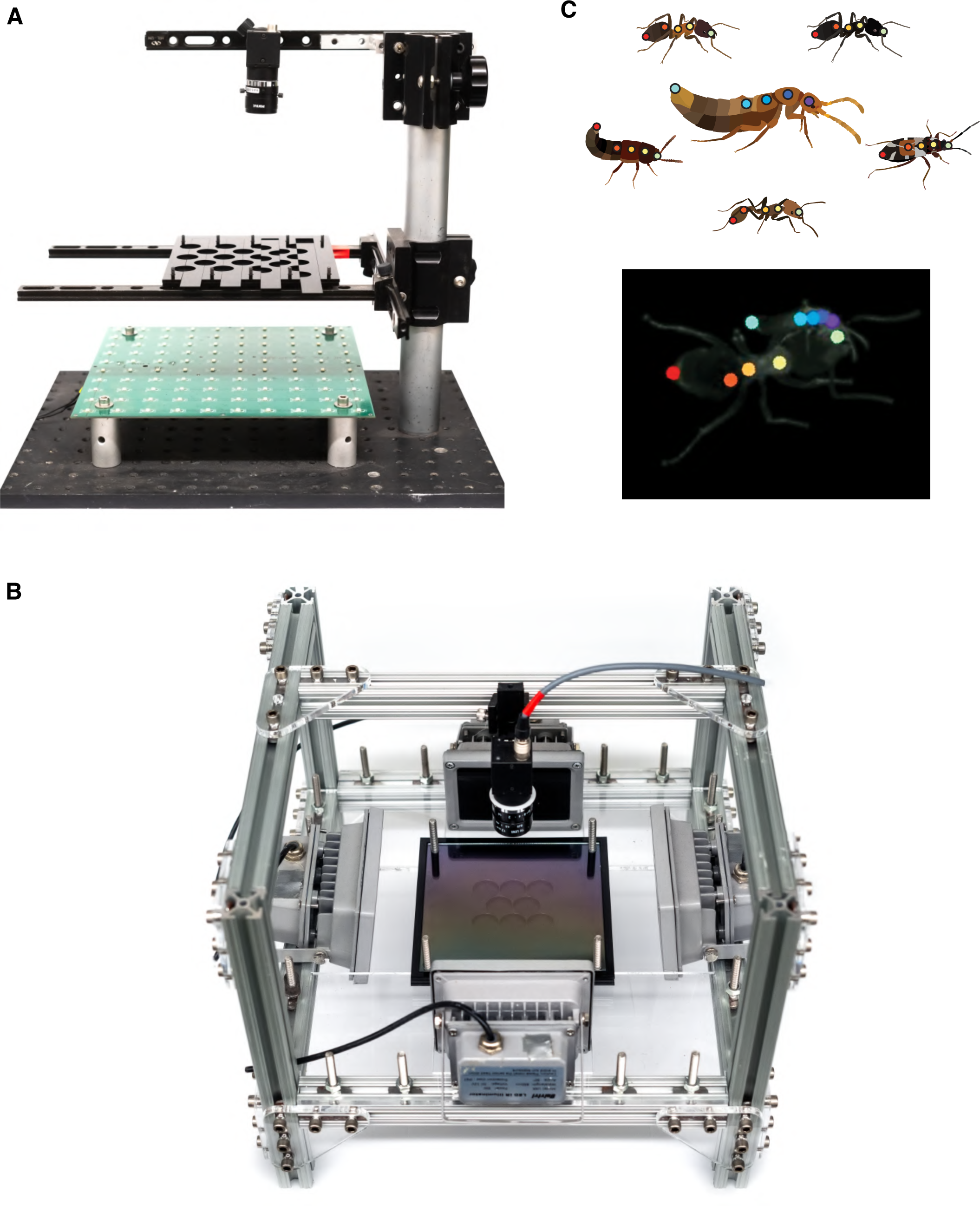
Behavioral arenas to probe grooming behavior. (**A**) Initial arena design for grooming assays. The arena is illuminated from below with IR LEDs and monitored from above with a Flir machine vision camera. The arena itself is composed of layers of IR transmitting acrylic, and all interactions occur in the dark inside a controlled temperature incubator. The arena wells are composed of half-circles; when loaded, the circles are staggered relative to each other to separate the animals, and one half can slide into place, allowing the animals to interact after they recover from cooling on ice, which is used to anesthetize them before loading them into the arena. (**B**) Second arena design tracking grooming behavior. The arena is illuminated from the sides with IR LEDs, and monitored from above with a camera. The arena is built with acrylic layers, but has no sliding mechanism after we found that keeping animals apart at the beginning of trials was unnecessary for most experiments. (**C**) A single DeepLabCut model was trained on annotated body positions of *Sceptobius* and other interacting insects. Shown are the key point locations used for the various interactors and for *S. lativentris*. Also shown is an example frame from a video of *Sceptobius* grooming an ant, showing keypoint locations.

**Figure S2.**
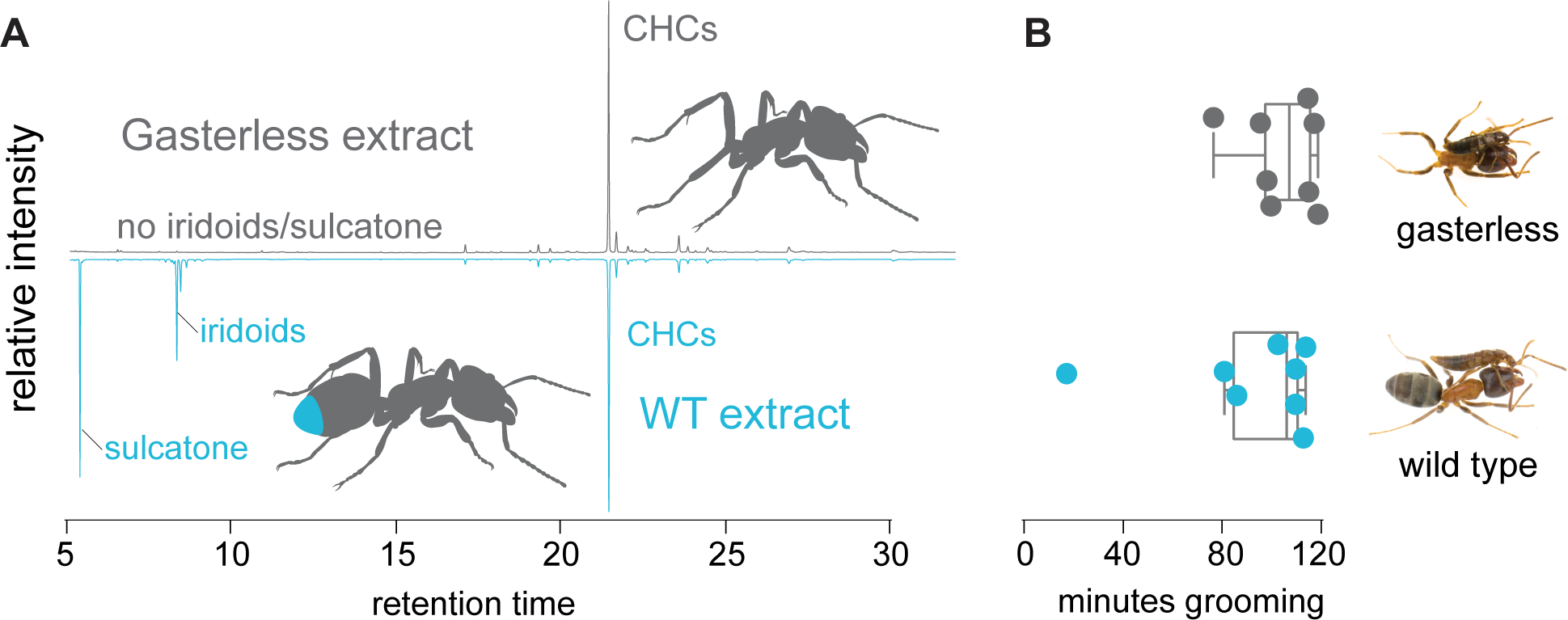
*Sceptobius* grooms gasterless (exclusively CHC-bearing) ants. (**A**) GC trace shows gasterless ants only bear CHCs, not iridoids, on their body surface. (**B**) *S. lativentris* grooms these ants with the same proclivity as it does intact ants, indicating that CHCs, and not other ant pheromones present in the gaster, release grooming behavior (blue region in cartoon ant abdomen indicates the pygidial gland—the source of iridoids).

**Figure S3.**
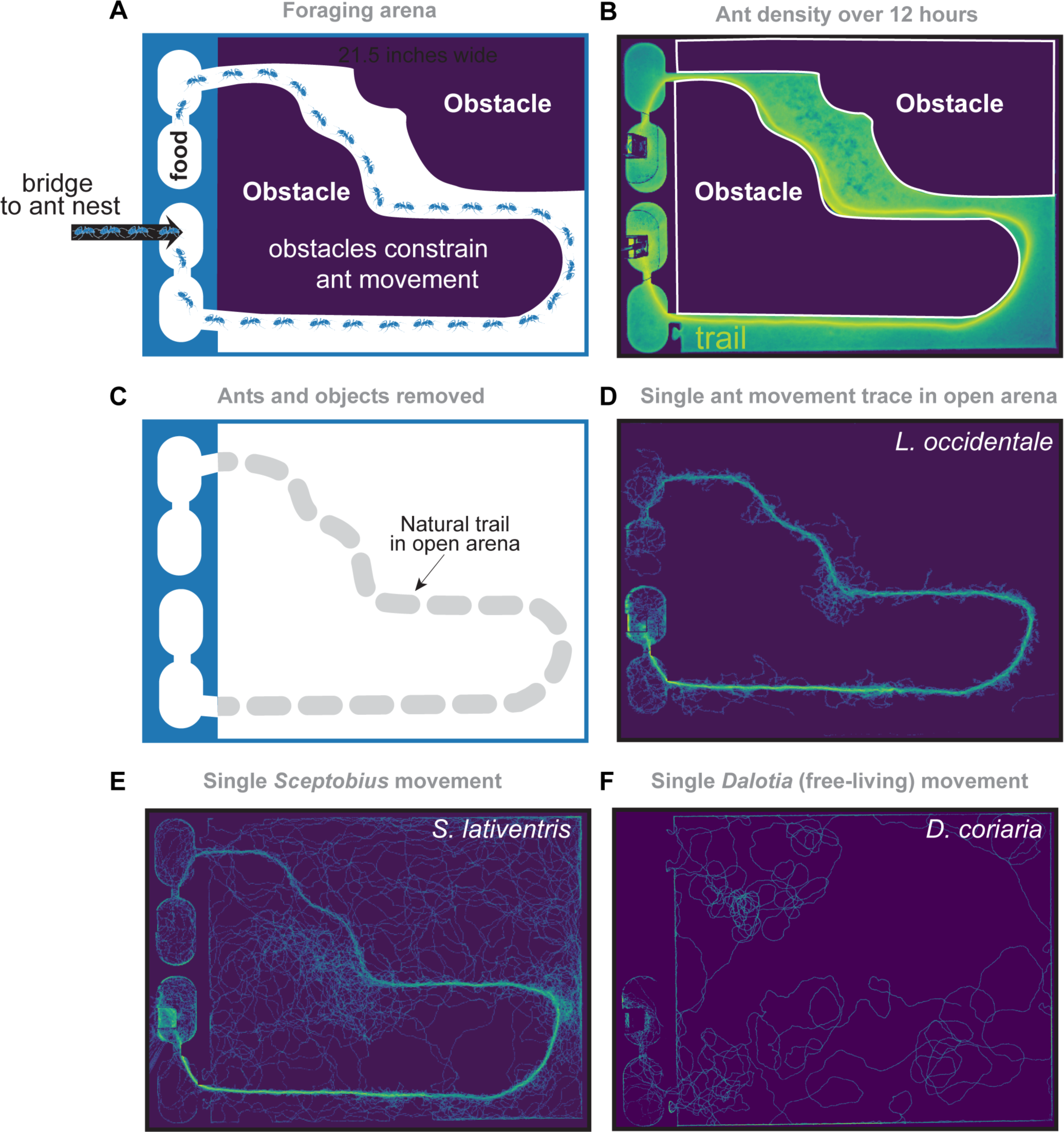
Assay design to probe natural trail following. (**A**) An *L. occidentale* ant colony was connected to the entrance of a behavioral arena with food positioned at the end of an obstacle-filled space. (**B**) Heatmap of the location of ant movement showing a path through the maze, indicating the location of the foraging trail formed by the ants. (**C**) After many hours, the ants and the obstacles were removed from the arena, leaving only a naturally laid chemical trail on the arena floor. (**D**) Single worker ants followed the predicted position of the trail with high accuracy. (**E**) Similarly, a single *S. lativentris* beetle closely followed the trail. (**F**) A non-symbiotic beetle, *Dalotia*, showed no trail-following behavior.

**Figure S4.**
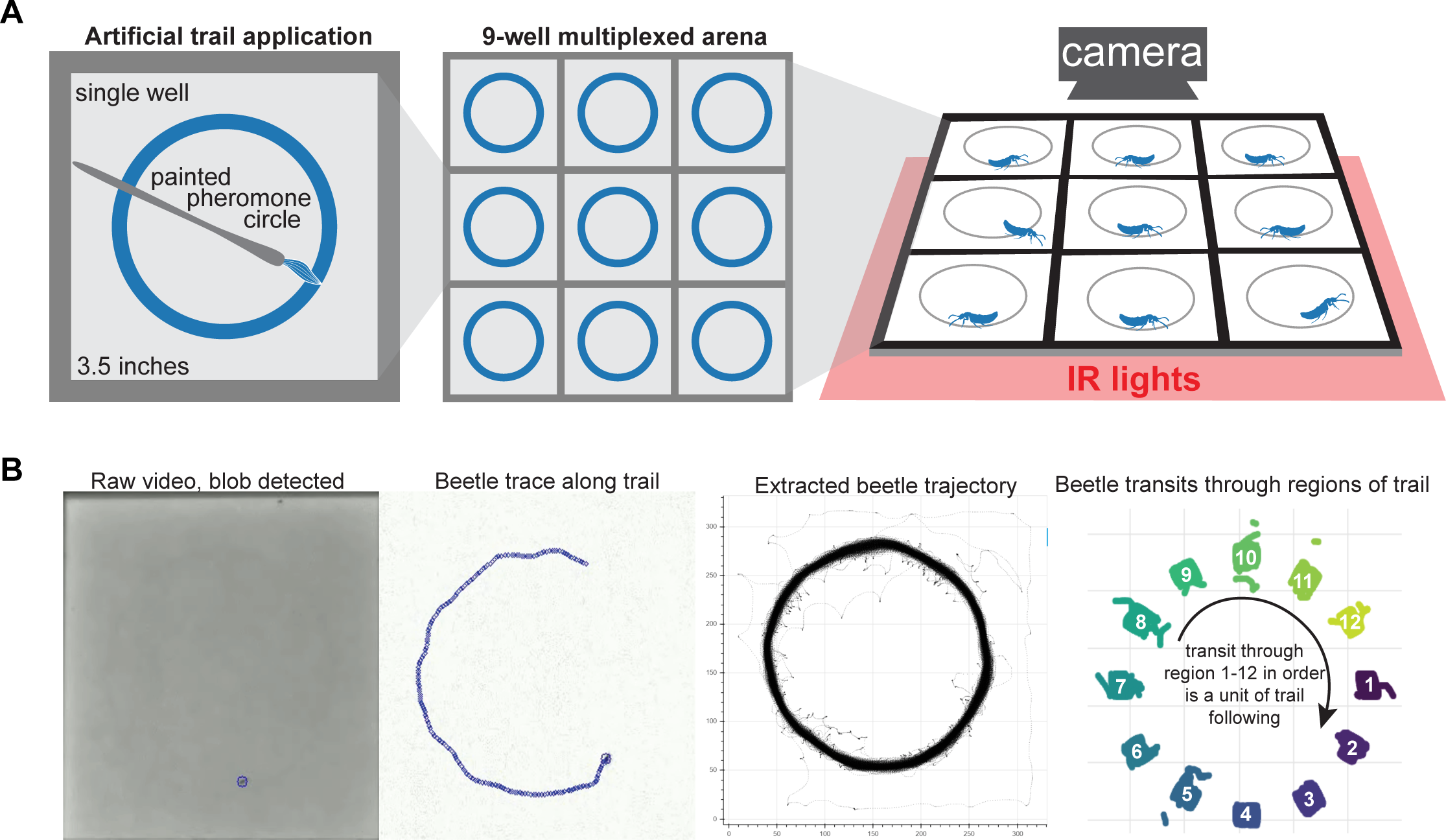
Assay to identify trail pheromones. (**A**) Different chemical components were painted in circles onto a ground glass surface in a multiplexed arena, and animal movement was tracked around the painted trails. (**B**) Blob detection on raw videos was performed after background subtraction. With the resulting trajectories, trail-following bouts were classified within the video when the animal transited though regions of interest along the trail in sequential order, without reversing direction or repeatedly transiting through a single section. This approach stringently and accurately provided the sections of the trajectory where the beetle walked in circular arcs around the arena.

**Figure S5.**
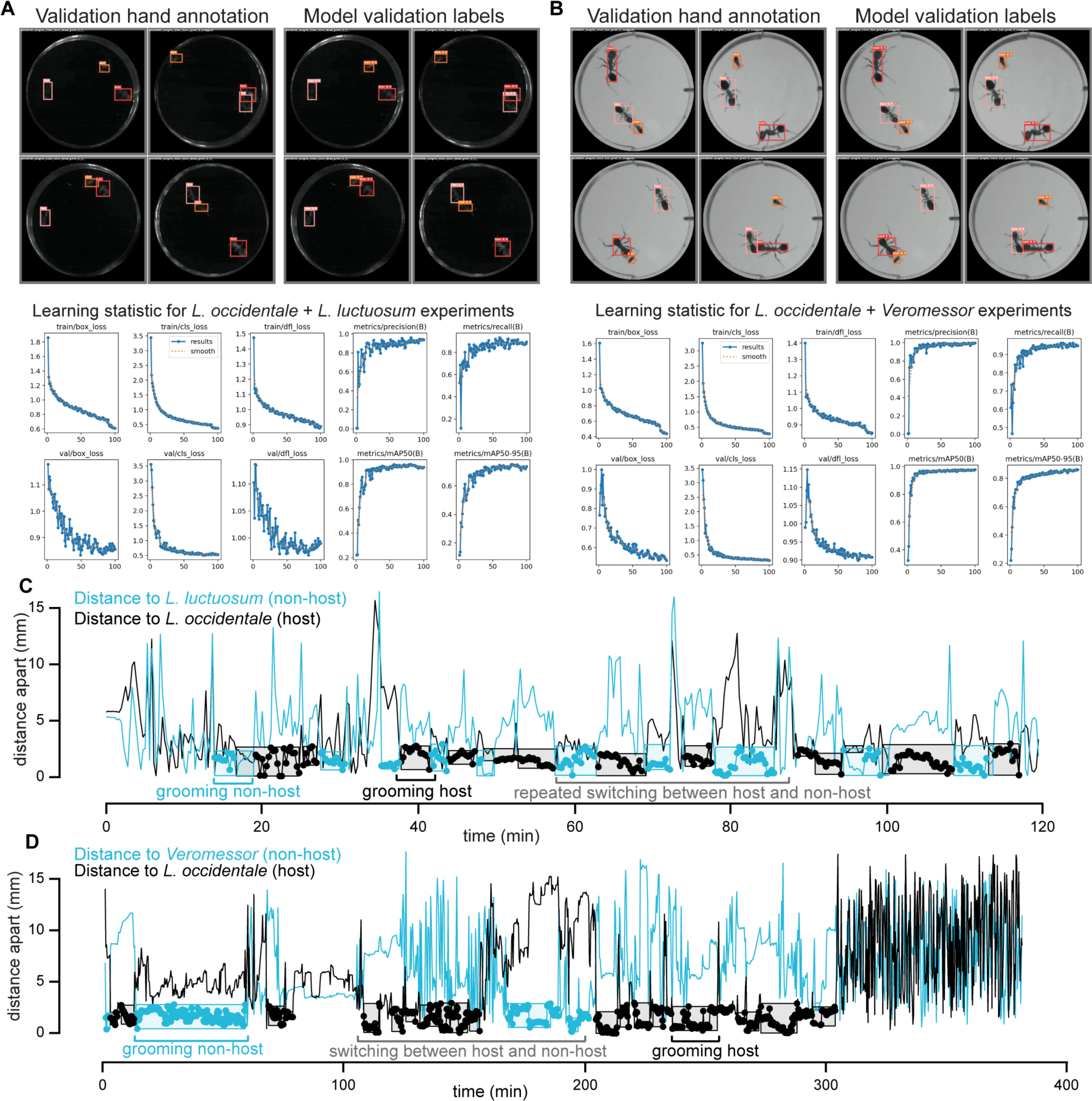
Machine vision performance and outputs for grooming preference assays, and example preference assay data. (**A**) Training data and training statistics for YOLO object detection models for preference assay analysis of *L. occidentale* vs *L. luctuosum*. Top panels show example bounding boxes for animals hand-annotated with CVAT, and bounding boxes generated by the trained model. Bottom panels show performance statistics of training the model. (**B**) Training data and training statistics for YOLO object detection models for preference assay analysis of *L. occidentale* vs *Veromessor*. Top panels show example bounding boxes for animals hand-annotated with CVAT, and bounding boxes generated by the trained model. Bottom panels show performance statistics of training the model. (**C, D**) Traces of the distance between animals during a behavioral trial show annotated grooming bouts on hosts and non-hosts: *L. occidentale* versus *L. luctuosum* (**C**), and *L. occidentale* versus *Veromessor* (**D**). Some trials show extensive switching between the host and non-host ants during the run, indicating a high proclivity to groom non-hosts even in the presence of hosts.

**Figure S6.**
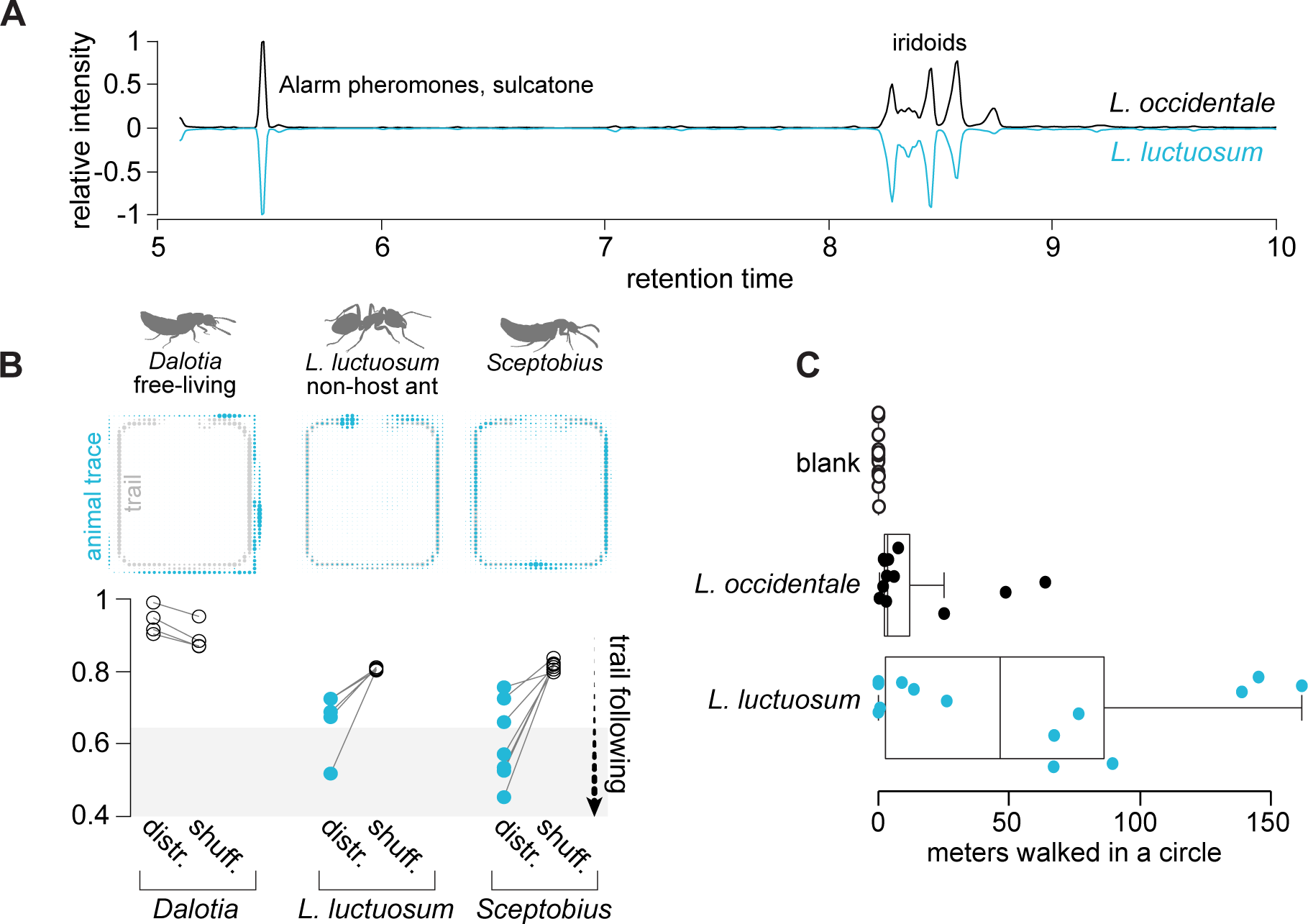
*Sceptobius* trail-following is not host specific. (**A**) GC trace of bulk extract of *L. occidentale* trail pheromones compared to its sister species, *L. luctuosum*. (**B**) Movement density plots show *S. lativentris* follows the trails of non-host sister ant species *L. luctuosum*. Ants also follow the trail, and free-living beetles do not. Using a dissimilarity measure derived from the Bhattacharyya distance, movement of *S. lativentris* and the ant correspond closely to the trail distribution (‘distr.’); randomly shuffling beetle movement traces abolishes the close match to the trail distribution, indicating that random movement cannot account for the correspondence of beetle movement with ant trail (‘shuff.’), whereas beetle movement in an empty arena had an equally negligible fit to trail shape as a randomly shuffled movement trace. (**C**) The number of meters that beetles followed ant extracts in a circle trail assay, showing extreme trail following with non-host compounds.

**Figure S7.**
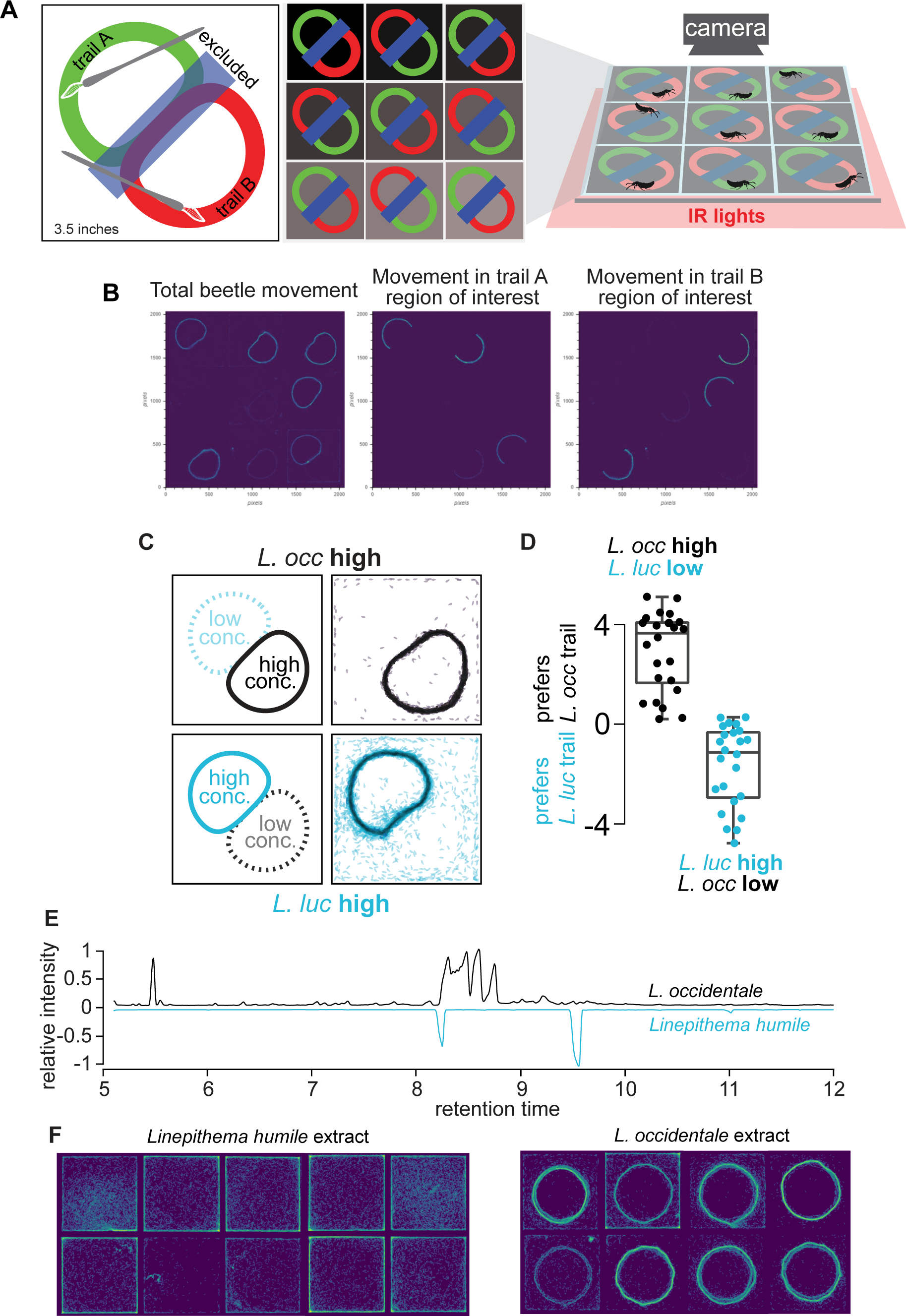
Analysis of multiplexed preference assay for trail chemicals. (**A**) Chemicals from different species were painted at different concentrations as semi-circular and abutting lobes. (**B**) Bulk movement of the beetle during the trial was calculated by subtracting subsequent frames from the behavioral trial. Manually labeled regions of interest (shown in **A**) were defined to extract the bulk beetle movement on the different arms of the trail (excluding the middle section where trails overlapped). For each trial, movements along a particular section of trail were summed, and these values were subtracted to obtain a metric for differential trail following of the two ant species (as in **D**). (**C**) For experiments, a section of the trial from one species was painted at high concentration. (**D**) Degree of following of trail corresponded with high concentration trails, regardless of which species chemicals these represented. (**E**) GC trace of crude *Linepithema humile* ant trail compounds compared to crude extract of *L. occidentale* trail chemicals, showing no overlap of peaks. (**F**) Beetles ignored *Linepithema humile* ant extracts painted in a circle in the multiplexed trail arena, but closely followed extracts of *L. occidentale*.

**Figure S8.**
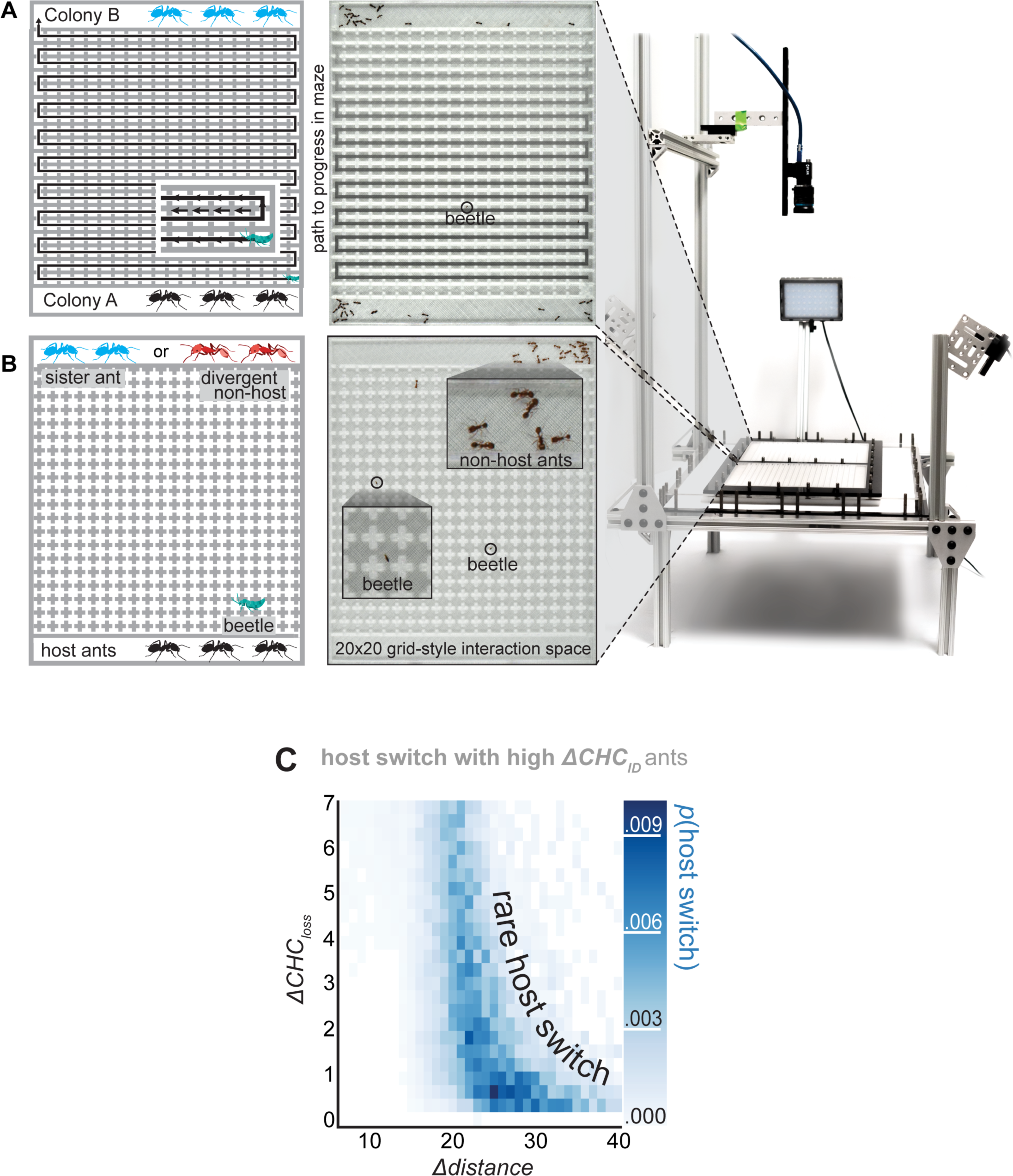
Cross-maze arena to probe dispersal abilities, and cross arena to probe host switching. (**A**) To test dispersal abilities of beetles, an arena was constructed with two flanking ant colonies (A and B), connecting by a zig-zagging corridor. This design created a maze with a minimal distance of ∼4 meters for beetles to cross from one colony to the other. Ants in both colonies were prevented from entering the arena by creating entrances from both colonies through which only *Sceptobius* could fit. (**B**) To observe beetles behaving in a space with two ant species, an arena was designed with a large grid of connected interaction chambers. Differing ratios of ants of host and non-host species, along with beetles, were introduced into colony areas flanking the central foraging arena. For setups in both **A** and **B**, LED photography lights were used to illuminate the arena from above on all sides. A wide-angle lens and high-resolution color camera were used to record behavior in the arena. (**C**) A rare additional case predicted by our in-silico model: if a beetle wanders away from ants long enough to lose almost all of its CHCs, yet retaining enough total CHC to prevent desiccation, it has a low but non-zero chance of host switching to even an aggressive non-host ant with high *ΔCHC_ID_*.

**File S1:**
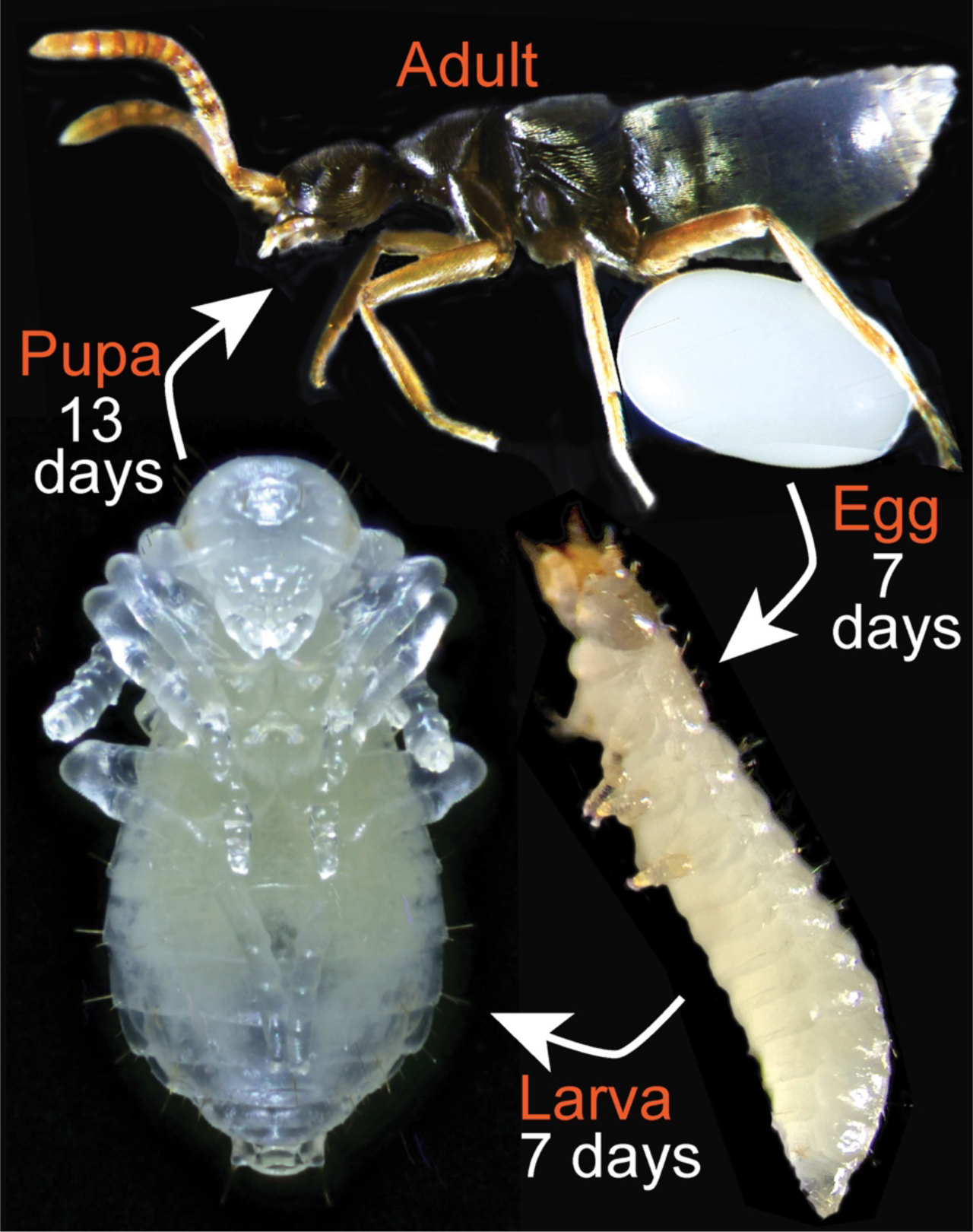
Life history of *Sceptobius lativentris*. The entire life cycle of *Sceptobius lativentris* takes places in close association with the nest of *Liometopum occidentale*. The host ant builds colonies inside trunks of coastal live oaks and bay trees, excavating the heartwood and building extensive labyrinthine carton nests (trabeculae). On acquiring a colony’s CHC profile via grooming, beetles gain access to the nest. Inside, beetles concentrate in large numbers inside brood galleries, where they feed on ant eggs and larvae. Adult beetles have also been reported to engage in oral trophallaxis with host workers (Danoff-Burg 1996). Reproduction occurs within the nest. Males and females are often observed copulating while simultaneously mounted onto worker ants. Imaginal development takes place at the nest boundary: females produce single, giant eggs that fill their entire abdomens, and oviposit into damp frass that has accumulated at the nest entrance from excavation of heartwood by the host ant. Larvae hatch from these eggs and remain secluded in the frass, away from ants, and do not apparently need to feed. Larvae rapidly progress to the pupal stage. Following pupal development, newly eclosed adults search for host workers, grooming them and re-integrating into the same, parental colony.

