## Supplementary material for "Enforced specificity of an animal symbiosis": File S1

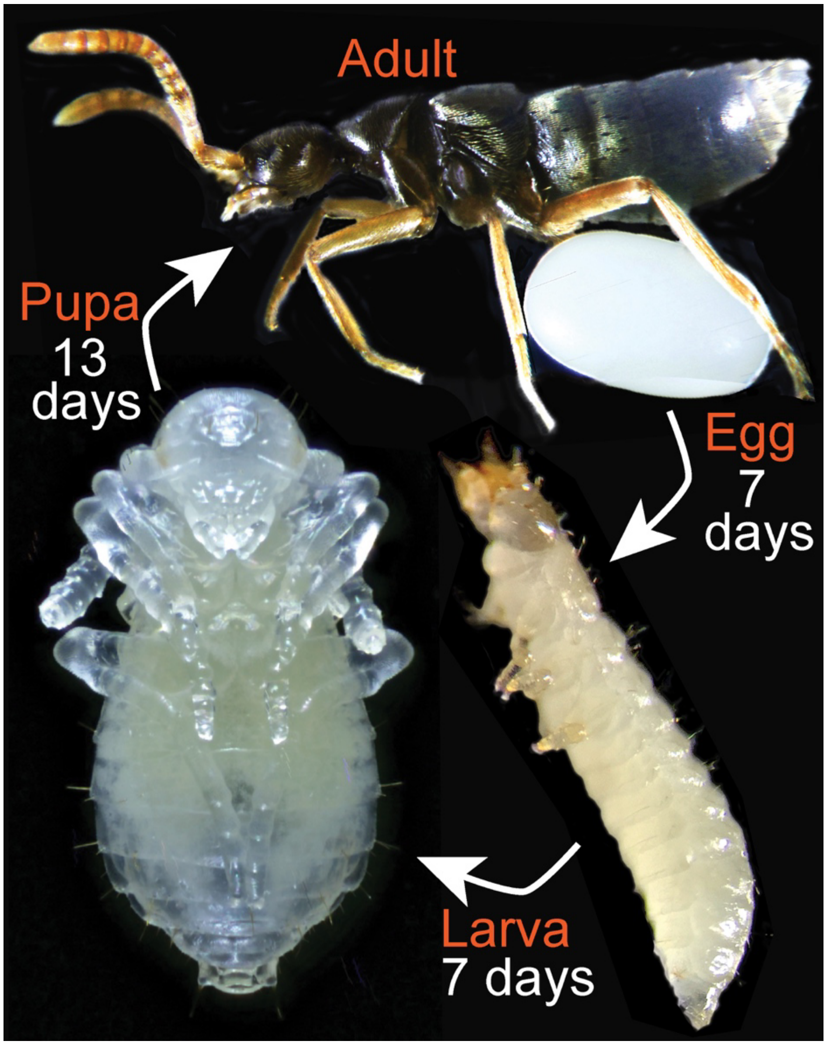


**File S1: Life history of *Sceptobius lativentris***

The entire life cycle of *Sceptobius lativentris* takes places in close association with the nest of *Liometopum occidentale*. The host ant builds colonies inside trunks of coastal live oaks and bay trees, excavating the heartwood and building extensive labyrinthine carton nests (trabeculae). On acquiring a colony's CHC profile via grooming, beetles gain access to the nest. Inside, beetles concentrate in large numbers inside brood galleries, where they feed on ant eggs and larvae. Adult beetles have also been reported to engage in oral trophallaxis with host workers (Danoff-Burg 1996). Reproduction occurs within the nest. Males and females are often observed copulating while simultaneously mounted onto worker ants. Imaginal development takes place at the nest boundary: females produce single, giant eggs that fill their entire abdomens, and oviposit into damp frass that has accumulated at the nest entrance from excavation of heartwood by the host ant. Larvae hatch from these eggs and remain secluded in the frass, away from ants, and do not apparently need to feed. Larvae rapidly progress to the pupal stage. Following pupal development, newly eclosed adults search for host workers, grooming them and re-integrating into the same, parental colony.
